# Parallel seed color adaptation during multiple domestication attempts of an ancient new world grain

**DOI:** 10.1101/547943

**Authors:** Markus G Stetter, Mireia Vidal-Villarejo, Karl J Schmid

## Abstract

Thousands of plants have been selected as crops, yet, only a few are fully domesticated. The lack of adaptation to agro-ecological environments of many crop plants with few characteristic domestication traits potentially has genetic causes. Here, we investigate the incomplete domestication of an ancient grain from the Americas, amaranth. Although three grain amaranth species have been cultivated as crop for millennia, all three lack key domestication traits. We sequenced 121 crop and wild individuals to investigate the genomic signature of repeated incomplete adaptation. Our analysis shows that grain amaranth has been domesticated three times from a single wild ancestor. One trait that has been selected during domestication in all three grain species is the seed color, which changed from dark seeds to white seeds. We were able to map the genetic control of the seed color adaptation to two genomic regions on chromosome 3 and 9, employing three independent mapping populations. Within the locus on chromosome 9, we identify a MYB-like transcription factor gene, a known regulator for seed color variation in other plant species. We identify a soft selective sweep in this genomic region in one of the crops species but not in the other two species. The demographic analysis of wild and domesticated amaranths revealed a population bottleneck predating the domestication of grain amaranth. Our results indicate that a reduced level of ancestral genetic variation did not prevent the selection of traits with a simple genetic architecture but may have limited the adaptation of complex domestication traits.

## Introduction

Crop domestication was the foundation for modern human societies and has taken place almost simultaneously in different regions of the world. Around 2,000 crops have been cultivated and underwent artificial selection during adaptation to human-mediated environments (Harlan 1992). This intense selection changed characteristic plant traits that are favorable for human cultivation (Olsen and Wendel 2013). Major crops, such as maize, rice, and wheat, combine multiple domestication traits, which together are referred to as domestication syndrome (Hammer 1984), reflecting their adaptation to agro-ecological systems and human preferences (Olsen and Wendel 2013; Li *et al*. 2006; Doebley 2004; Peleg *et al*. 2011). The domestication syndrome includes production-related traits, like reduction of seed shattering, larger seed size, loss of seed dormancy, and traits that reflect human preference, including taste changes, reduced toxin content, and seed or fruit color variation (Hammer 1984; Lenser and Theißen 2013).

Crop domestication is a long and fluid process rather than an instantaneous event, and for this reason crops differ in their degree of domestication (Stetter *et al*. 2017a; Gaut *et al*. 2018). Plants with different degrees of domestication have been cultivated over similar time periods (MacNeish 1965), but fully domesticated crops that combine multiple domestication traits are few in comparison to hundreds of cultivated crops with a weak domestication syndrome (Meyer *et al*. 2012). A weak domestication syndrome cannot be explained by weak selection during domestication, given a long period of cultivation and the central importance of numerous minor crops for early agricultural societies. Alternatively, a weak domestication syndrome may reflect genetic constraints or missing functional genetic variation in the wild ancestors (Walsh and Blows 2009). Incompletely domesticated crops are therefore suitable models to investigate the role of genetic constraints and missing diversity, because domestication traits under selection are known and multiple incompletely domesticated species are available as replicates (Purugganan and Fuller 2009; Purugganan 2019).

The pseudo-cereal grain amaranth is an example of a crop with a long cultivation history, but a weakly pronounced domestication syndrome. Traits like reduced spines on inflorescences and increased seed number per inflorescence reflect domestication-related selection, but other typical domestication traits of grain crops, such as increased seed size and the loss of seed shattering are missing and therefore limit the yield potential of grain amaranth (Sauer 1967; Brenner *et al*. 2010; Stetter *et al*. 2017b). One conspicuous domestication trait of cultivated amaranth is a change in seed color, as all wild amaranths have dark seeds, whereas most cultivated grain amaranth landraces have white seeds (Stetter *et al*. 2017b). The reasons for selection of white seed color remain unknown, but may be linked to agronomic traits like faster seed germination or improved seed processing by milling (Lenser and Theißen 2013).

Three species of *Amaranthus* are cultivated as grain crops in South (*Amaranthus caudatus* L.) and Central America (*A. hypochondriacus* L. and *A. cruentus* L.) (Brenner *et al*. 2010). They are closely related and were likely domesticated from the same wild ancestor, *A. hybridus L*., but it remains unclear if three separate or a single domestication process gave rise to the three grain amaranth species (Sauer 1967; Kietlinski *et al*. 2014; Stetter and Schmid 2017). The taxonomic differentiation of amaranth species is based on morphological traits, in particular on the appearance flower and inflorescence morphology (Sauer 1967). Archaeological findings show that amaranth seeds were collected already 8,000 years ago in South America (Arreguez *et al*. 2013) and 6,000 years ago in North America (MacNeish 1965). Historical accounts suggest that grain amaranths were of similar importance for the agriculture of pre-Colombian societies as maize (Sauer 1967), but amaranth cultivation rapidly declined after the Spanish conquest of the Americas. Nowadays, amaranth grains are gaining importance for human nutrition, due to their high concentration of essential amino acids, which provide high nutritional value (Downton 1973; Rastogi and Shukla 2013) and low gluten content that make it a suitable cereal replacement for patients with celiac disease (Ballabio *et al*. 2011).

In this study, we investigate the domestication process of amaranth by combining whole genome resequencing of a diverse set of 121 genebank accessions of the three grain amaranth species and two wild relatives, with an F_2_ mapping population from a cross of domesticated and wild individuals, and two bulk populations from advanced breeding material segregating for seed color. To characterize seed color as exemplary domestication trait, we combine genetic and evolutionary analyses to infer the role of selection and genetic constraints during domestication. Our results provide strong evidence that grain amaranth was domesticated three times and that white seed color was independently selected in each grain species. We identify a MYB-like transcription factor gene within a QTL region as potential regulator of seed coat color variation of grain amaranth. Based on a low ancestral effective population size, we suggest that the domestication of grain amaranth was limited by the lack of functional diversity for complex domestication traits at the time of domestication.

## Results and discussion

### Resequencing and variant characterization

We resequenced a diverse set of 121 genebank accessions that represent the native distribution ranges of grain amaranth. The set included the three grain amaranths, caudatus (*A. caudatus* L., *n* = 32), cruentus (*A. cruentus* L. *n* = 25), and hypochondriacus (*A. hypochondriacus, n* = 24), the two wild relatives hybridus (*A. hybridus* L., *n* = 19) and quitensis (*A. quitensis* (Kunth), *n* = 19), and two individuals that were labeled as ‘hybrids’ or *A. powellii* L. in their passport data (Table S1). The average sequence coverage per accession was 9x and 94% of reads could be mapped to the hypochondriacus 2.1 reference genome (Lightfoot *et al*. 2017). There was no difference in the proportion of mapped reads between species indicating an absence of a mapping bias in the non-hypochondriacus populations (Table S2). After variant calling, we identified over 29 million biallelic variants with less than 30% missing data and from these we extracted a final data set of 17,157,381 million biallelic SNPs for further analysis.

### Three domestication attempts from a single common ancestor

In numerous crop species, domestication led to a reduction in genetic diversity due to population bottlenecks and strong directional selection (Hufford *et al*. 2012; Stein *et al*. 2018) Nucleotide diversity (*π*) in all three grain amaranths (caudatus: 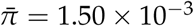; cruentus: 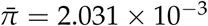; hypochondriacus: 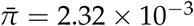) was reduced by a factor of two to three relative to hybridus 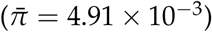, although the wild relative quitensis 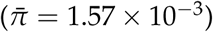 has similar values as the cultivated amaranths (Figure 1A). Signals of Tajima’s *D* were also similar across species and indicate recent population growth, with negative values for *D*. Average Tajima’s *D* was lowest for quitensis and close to zero for caudatus (Figure 1B).

**Figure 1.**
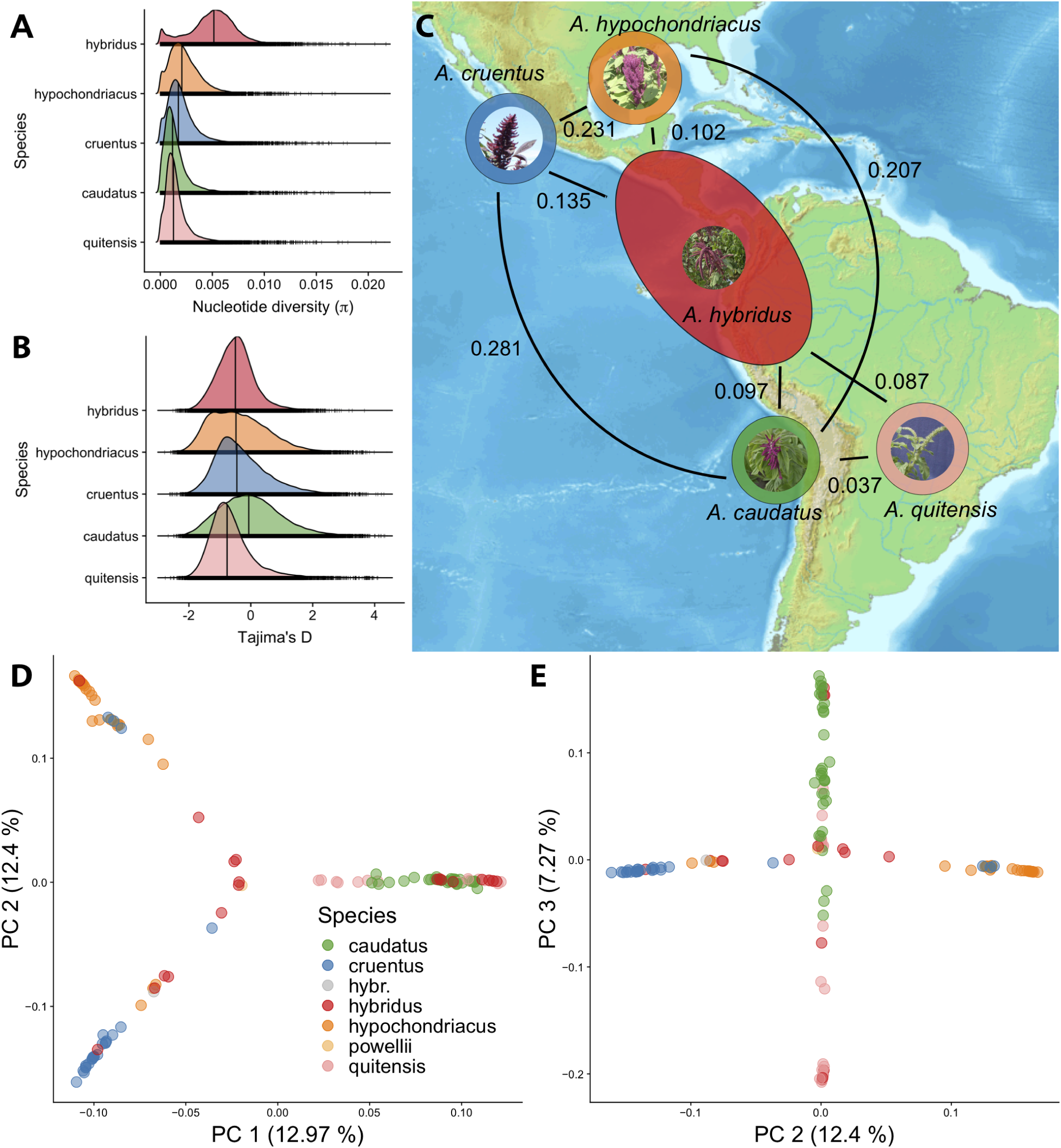
Distribution and diversity of amaranth. **A)** Nucleotide diversity (*π*) distribution of 10 kb windows, vertical line denotes mean. **B)** Distribution of Tajima’s D in 10 kb windows, vertical line denotes mean. **C)** Circles show the approximate geographic distribution of each species (The distribution of hybridus overlaps with all three grain species). Values on the lines show genome wide pairwise *F*_*st*_ comparisons between species. **D)** PCA of 121 individuals that indicates the differentiation between Central and South American accessions on PC 1, and between the grain amaranths from Central America on PC 2. **E)** shows the differentiation between caudatus and quitensis on PC 3. Coloring follows the description in (D). The grain amaranths caudatus, cruentus and hypochondriacus are shown in green, blue and orange, respectively, and the wild ancestors hybridus, quitensis in red and light red.

Using the fixation index, *F*_*st*_, as measure of genetic differentiation we tested different hypotheses of domestication. If all three grain amaranth species were domesticated in a single process, the level of genetic differentiation should be lower between the three domesticated than between the wild and domesticated amaranths. A single domestication was not supported by our data, because the three domesticated amaranths were more strongly differentiated from each other 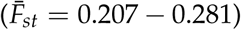 than each domesticated amaranth from their wild ancestor, hybridus (0.097 – 0.135; Figure 1C).

This result is consistent with the analysis of genetic population structure. An ADMIXTURE analysis revealed between three and seven clusters, which separate the three grain amaranths but cluster wild relatives with domesticated accessions from the same geographic regions (Figure S1). A similar pattern was found in a Neighbor Joining tree (Figure S3). A principal component analysis (PCA) clusters accessions into three groups (Figures 1 and S4). The first principal component (PC) separated all wild and domesticated South American accessions (caudatus, quitensis, and hybridus) from all Central American accessions (cruentus, hypochondriacus, and hybridus). The second PC differentiated the two Central American grain species, and the third PC separated the South American hybridus and quitensis from the domesticated caudatus accessions. We obtained a similar population structure with a Community Oriented Network Estimation analysis (CONE; Kuismin *et al*. 2017), which differentiated all accessions into three sub-networks that separate the grain amaranths with no edges between them (Figure S5). Taken together, the different analyses of population structure suggest three domestications, because each of the three grain amaranth species clustered with a subset of genetically differentiated hybridus accessions that originate from the same geographic region as their genetically most similar cultivated amaranths. An alternative model of consecutive domestications after an initial domestication of an ancestral grain amaranth from hybridus would have resulted in a closer relationship between caudatus, cruentus and hypochondriacus than to the wild ancestor hybridus, which was not observed in our data.

Previous studies observed a very close relationship between the wild quitensis and domesticated caudatus, which are sympatric in South America. This led to the hypothesis that quitensis is the direct ancestor of caudatus (Sauer 1967; Kietlinski *et al*. 2014). Our analysis of *F*_*ST*_-values also showed a weaker differentiation of caudatus from quitensis (0.037) than from hybridus (0.097; Figure 1C), but hybridus is geographically structured (Figure S1) and South American hybridus is more closely related to caudatus and quitensis than caudatus is to quitensis (Figure 1, S4 and S1). In earlier work with a larger sample of South American amaranths we have shown a close relationship between hybridus and quitensis and suggested hybridus as ancestor of caudatus (Stetter *et al*. 2017b). Taken together, the inferred population structure was highly consistent with a model of repeated domestication of each of the three grain amaranths from geographically separated and genetically differentiated populations of hybridus.

### Common major domestication QTL in all three domesticated amaranths

We next investigated the relationship between the genetic history of the grain amaranths and seed color, which is a domestication trait in South American caudatus (Stetter *et al*. 2017b) and therefore tested whether white seeds are also a derived domestication trait in cruentus and hypochondriacus. In our diversity panel, all accessions of the wild ancestors have dark seeds, whereas the majority (73 to 82 %) of the grain amaranth accessions have white seeds (Figure S6). Since the population genetic analysis suggests multiple domestication attempts for each grain amaranth, we investigated whether they share the genetic basis of white seed color and if this was selected independently.

A classical genetic analysis based on the segregation ratio in crosses between white and dark seeded amaranths suggested that seed color is controlled by two genetic loci and white seed color is recessive (Kulakow *et al*. 1985). To identify the genes that control seed color, we conducted a QTL analysis in a segregating *F*_2_ population of a cross between a cultivated caudatus (white seeds) and wild quitensis (dark seeds) and a bulked segregant analysis of a population consisting of a diverse panel of cruentus and hypochondriacus individuals. We also carried out a genome wide association study (GWAS) with 116 accessions from the five species of the diversity panel and a reference-free association study of *k*-mers performed in each of the three different genetic groups detected with CONE and supported by the PCA. We identified the same two major QTL on chromosomes 3 and 9 in all three populations (Figures 2 and S7). The QTL on chromosome 3 was located in a narrow genomic interval of 100 kb in the GWAS. The QTL on chromosome 9 extended over a larger interval in the F_2_ mapping and bulk segregant analysis. In the GWAS, SNP markers associated with seed color were also distributed over a large region of ca. 600 kb (Figure S7). Such a large interval is not expected in a GWAS on a genetically diverse panel consisting of two wild and three domesticated amaranth species, even after correction for population structure. The large region on chromosome 9 that we observed in all three mapping populations could be caused by suppression of recombination in this region between color morphs. A potential reason for this signal could be a chromosomal inversion during domestication (Griffiths *et al*. 2005). We compared linkage disequilibrium (LD) between white and dark seeded groups in each species, which indicated high LD in white cruentus and hypochondriacus, but not in caudatus, suggesting that the large QTL region was mainly caused by Central American amaranth (Figure S8). In addition, an alignment of chromosome-wide CDS regions between the hypochondriacus reference with white seeds, and *A. tuberculatus* (Kreiner *et al*. 2019), a wild relative with dark seeds, showed an inverted segment between the two genomes that partly included the QTL region. In addition, the alignment showed a duplication of parts of the region in *A. tuberculatus*. While this pairwise comparison does not allow to infer the origin of the inversion or its segregation in the crop species, it does suggest the presence of a structural variant in hypochondriacus which could have caused the observed GWAS result.

**Figure 2.**
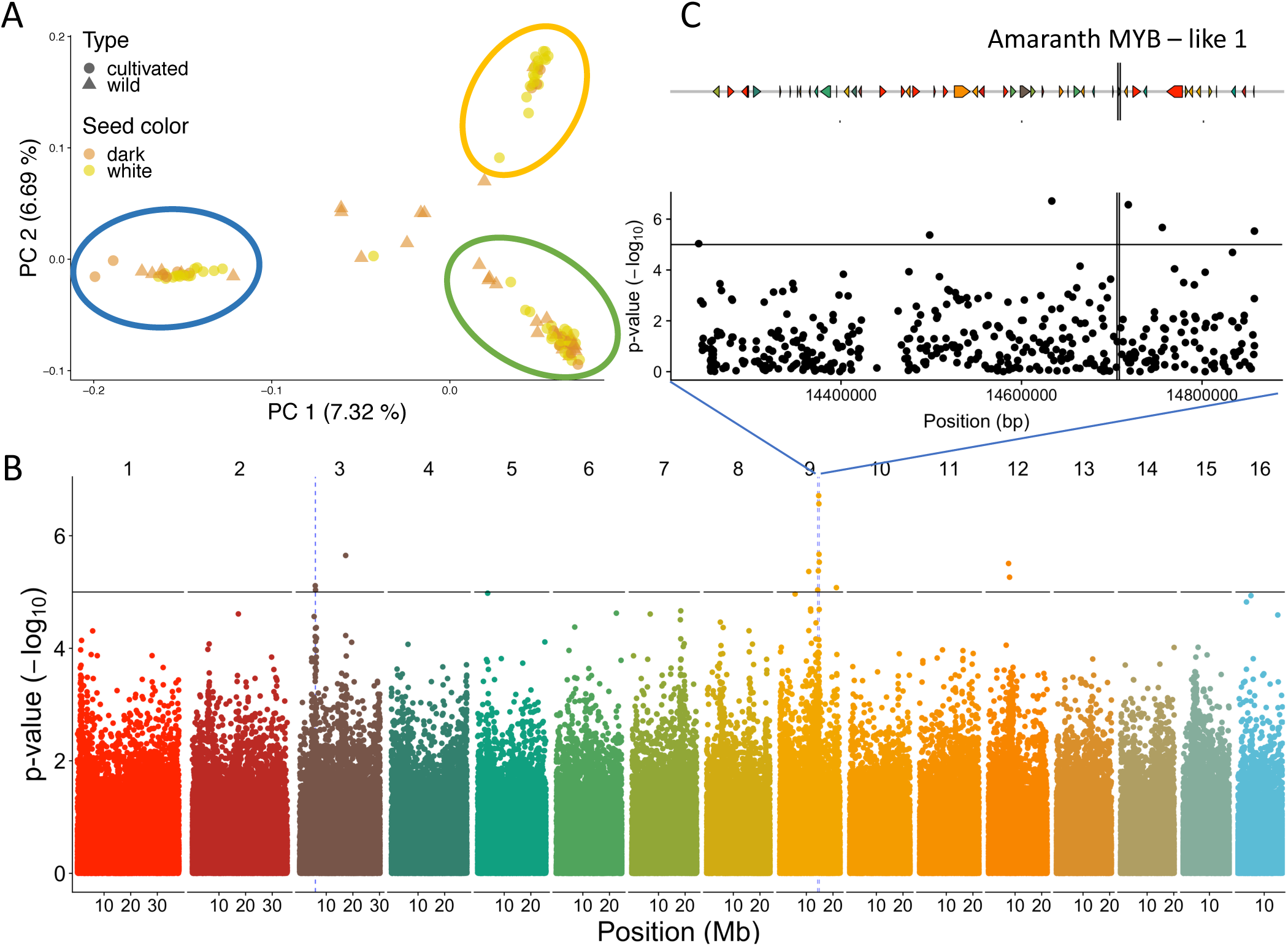
Genetic control of seed color variation **A)** Local PCA of QTL regions, using only SNPs within QTL regions for seed color as indicated by blue lines in B. Circles indicate the predominant crop species in the cluster, cruentus (blue), caudatus (green) and hypochondriacus (orange). **B)** Genome wide association mapping within 116 diverse amaranth accessions from the two wild amaranths and the three grain amaranth species. The horizontal line indicates a significance level of 10^−5^. Dashed blue lines indicate QTL regions for seed color as identified in three independent populations. **C)** Zoom of QTL region on chromosome 9 and genes within this region. The candidate gene AmMYBL1 is marked by vertical lines.

We further tested whether white seed color originated once or multiple times during domestication by conducting a local PCA on the two QTL regions in the diverse GWAS panel (Figure 2A and S10). A single evolutionary origin of white seeds is expected to separate white and dark seeded accessions into two separate groups independent of the species. The analysis revealed three groups that corresponded to the three grain amaranth species, in which the wild ancestors were nested in each group, similar to the genome-wide population structure, showing that a conversion from dark to white seeds was selected independently in each of the three grain amaranths (Figures 2A and S10).

### MYB-like gene as candidate for convergent seed color conversion

To identify candidate genes controlling seed color, we investigated the annotation of genomic regions with significant SNPs in the GWAS. The QTL region on chromosome 3 harbored five genes and the large region on chromosome 9 included 50 genes (Table S3). The latter region included a MYB-like transcription factor gene. The MYB transcription factor family is widely known for its key regulatory function in color phenotypes across the plant kingdom (Gates *et al*. 2016) and their role in color adaptation during crop domestication has been shown for several grain crops (Paauw *et al*. 2019). The MYB gene on chromosome 9 is an ortholog of the Anthocyanin Regulatory C1 gene in maize (Cone *et al*. 1986), and the MYB2 gene in grape (Walker *et al*. 2007). The latter is responsible for red and green grape skin and the change from red to green grapes is caused by the insertion of a transposable element (TE) within the gene (Walker *et al*. 2007). In blood orange, an expression change of a MYB gene increases pigmentation in the flesh and leads to the dark red flesh color (Butelli *et al*. 2012). In beet, a close relative of amaranth, a MYB transcription factor has been shown to control leaf pigmentation (Hatlestad *et al*. 2015). This gene has also been shown to be functional in amaranth shoots, confirming the general contribution of MYB transcription factors in amaranth (Hatlestad *et al*. 2015).

Despite the large number of genes in this region, the MYB transcription factor (Amaranth MYB-like 1, short: AmMYBL1) is a promising candidate gene for seed coat color. The sorted haplotype representation of the AmMYBL1 region shows that the three grain amaranths are differentiated in this region and that individuals with dark seeds have haplotypes more similar to the wild relatives than white-seeded grain amaranths (Figure S11A).

To identify potential insertions or deletions responsible for the change in seed color, we used a reference free *k*-mer based association method (HAWK; Rahman *et al*. 2018). HAWK identified *k*-mers that were significantly associated with white seed color among the South American accessions, but not among the Central American groups of hypochondriacus and cruentus. We *de novo* assembled the significant k-mers and mapped the resulting contigs to the amaranth reference genome and found that three of the four assembled contigs locate to the QTL region on chromosome 9. One contig mapped 2.3 kb downstream of the candidate MYB gene, and the other two 120 kb and 136 kb upstream of candidate MYB-like transcription factor gene.

In addition, we annotated the unfiltered variants segregating within gene and in the 2kb upstream and downstream region of the candidate gene to identify putative functional variants. Within this region 9 missense mutations were segregating in our diversity panel accessions (Figure S12). In summary, the k-mer-based mapping and functional annotation of polymorphisms are consistent with a role of the candidate MYB gene in a white seed color, but further investigations are required to identify the causal variants.

Since MYB transcription factors were shown to form a protein complex with bHLH and WD-repeat transcription factors (Ramsay and Glover 2005), we searched for genes encoding such proteins in the QTL regions on chromosome 3 and 9. Using sequence similarity to ortholog sequences from other plants we were not able to identify such genes in these two intervals suggesting that other genes in the QTL region on chromosome 3 influence seed color (Jun *et al*. 2015).

### Genome wide signatures of selection

The separate domestication of the three grain amaranths suggests that their genomes exhibit independent but potentially convergent domestication-related footprints of selection in their genomes. In particular, we expected to identify selective sweeps of independent *de novo* mutations in the genomic regions identified in the QTL analysis for white seed color because this trait was selected independently during each domestication. In major crops hard selective sweeps at domestication genes have been identified (Tian *et al*. 2009; Olsen *et al*. 2006). To detect hard selective sweeps and to differentiate them from soft selective sweeps, we used three different selection tests. We first compared the overlap of selective sweep locations identified by SweeD (Pavlidis *et al*. 2013) in each of the five amaranth species to test whether selection affected the same or different genomic regions. Of the top 0.5 % (99 total) outlier regions, 17 overlapped in the two wild species quitensis and hybridus (*p* ≤ 7.073 × 10^−22^). The overlap of putative sweep regions was lower between wild and domesticated amaranths, but was also significant (4 of 99, *p* ≤ 0.002). In contrast, outlier regions did not overlap between the three domesticated amaranths, and were unique to each species (Figure 3), suggesting that selection during the three domestication processes affected different genes. We did not identify hard selective sweeps in the two genomic regions on chromosomes 3 and 9 that harbor the seed color QTLs in any of the three domesticated amaranths with both SweeD (Figure S13) and RAiSD (Figure S15). To test whether soft selective sweeps my have occurred in the seed color QTL regions, we used the G2/G1 test (Harris *et al*. 2018). This method identified a selective sweep in hypochondriacus within the QTL region on chromosome 9. Two of the ten significant outliers on this chromosome were located in within the QTL region (Figure S16). One of these regions had the highest G123 value of 0.4 indicating a soft selective sweep. A comparison of the ten most significant outliers identified by SweeD and RAiSD, which both detect only hard sweeps to outliers identified by G2/G1, which is able to detect both hard and soft selective sweeps, provided further evidence for a soft selective sweep during the rise of white seed color in hypochondriacus (Figure S16). Such a result may be either explained by multiple new mutations in the causative region with a species during domestication, or selection on standing variation. Standing genetic variation for the recessive white seed color could persist at low frequency in the wild, despite potential selection against white seeds. A frequency of up to 27 % dark seeded accessions in grain amaranths further suggests the absence of a complete hard selective sweep during domestication.

**Figure 3.**
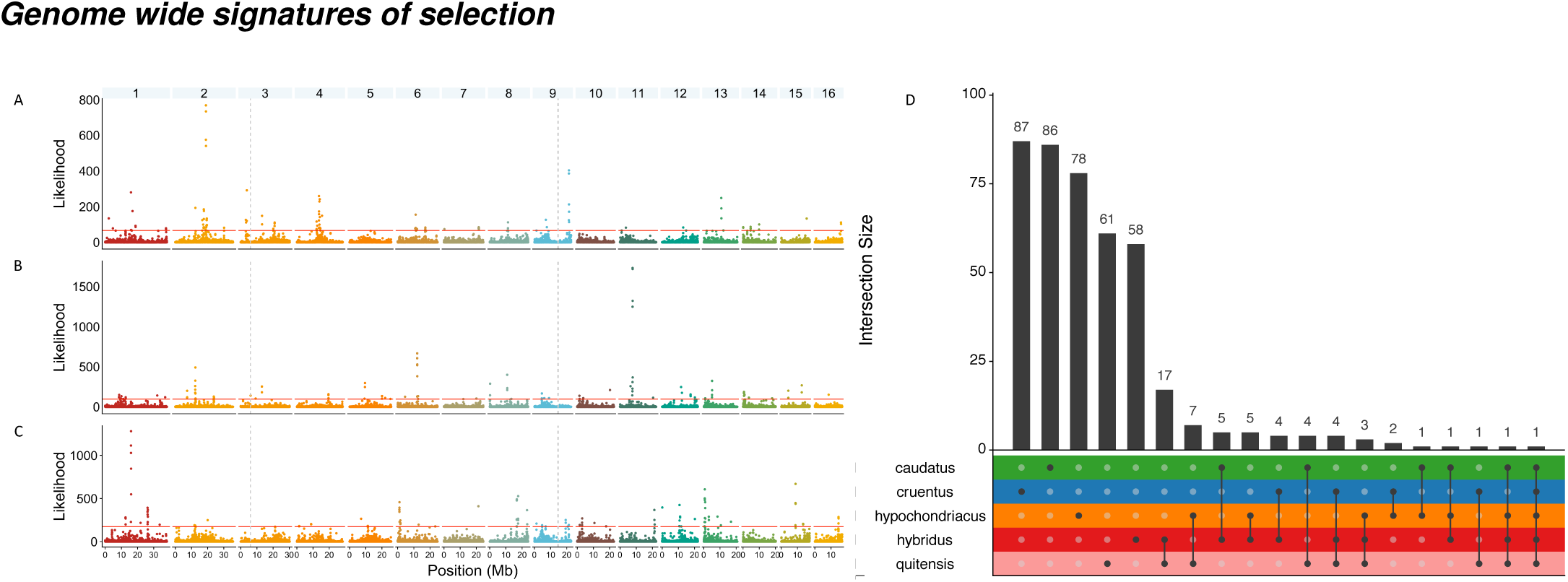
Detection of selection with SweeD in the three grain amaranths. Selective sweeps along the genome in cultivated grain amaranths in **A)** caudatus, **B)** cruentus and **C)** hypochondriacus. The horizontal line shows the top 0.05 % likelihood scores within each species **D)** Overlap between sweep regions of the top 0.5% outlier regions in amaranth species (three crop species and two wild relatives). Lines between species show comparisons and bar height represents the number of overlaps.

### Population history and gene flow between wild and domesticated amaranths

Our analysis of population structure did not indicate a strong isolation between wild relatives and domesticated amaranths, which may reflect extensive gene flow. In addition, the incomplete fixation of white seed color in each of the three grain amaranths and the absence of strong signals of selection may reflect ongoing gene flow from wild into domesticated amaranths. It is well established that crops and their wild relatives intercross in the wild and exchange adaptive traits (Janzen *et al*. 2018). For example, adaptive introgression from wild relatives was observed in major crops like maize (Hufford *et al*. 2013), barley (Poets *et al*. 2015), rice (Cubry *et al*. 2018; Meyer *et al*. 2016) and pearl millet (Burgarella *et al*. 2018). To characterize the extent and directions of gene flow in amaranths, we generated a population tree and estimated gene flow with TreeMix (Figure 4). The resulting unrooted tree is consistent with the population structure analysis, but also indicates a substantial level of gene flow between different groups. All three grain amaranths show traces of gene flow from or into wild relatives, and between Central and South American amaranths indicating a history of migration and exchange of genetic variation. Such patterns of gene flow would allow the introgession of white seed color genetic variants on chromosome 3 and 9 into all three grain amaranths after initial selection in one population. However, both local PCA (Figure S10) and haplotype structure analyses (Figure S10) at the two QTL regions and the AmMYBl1 gene did not cluster genetic variation by seed color as expected under model of a single origin, but by the overall population structure. Our results therefore suggest that white seeds originated and were selected independently in each grain amaranth and did not result from a single domestication or post-domestication introgression. However, ongoing gene flow from wild into domesticated amaranths may have prevented the fixation of white seed color alleles in the grain amaranths, as we proposed previously for the South American caudatus (Stetter *et al*. 2017b).

**Figure 4.**
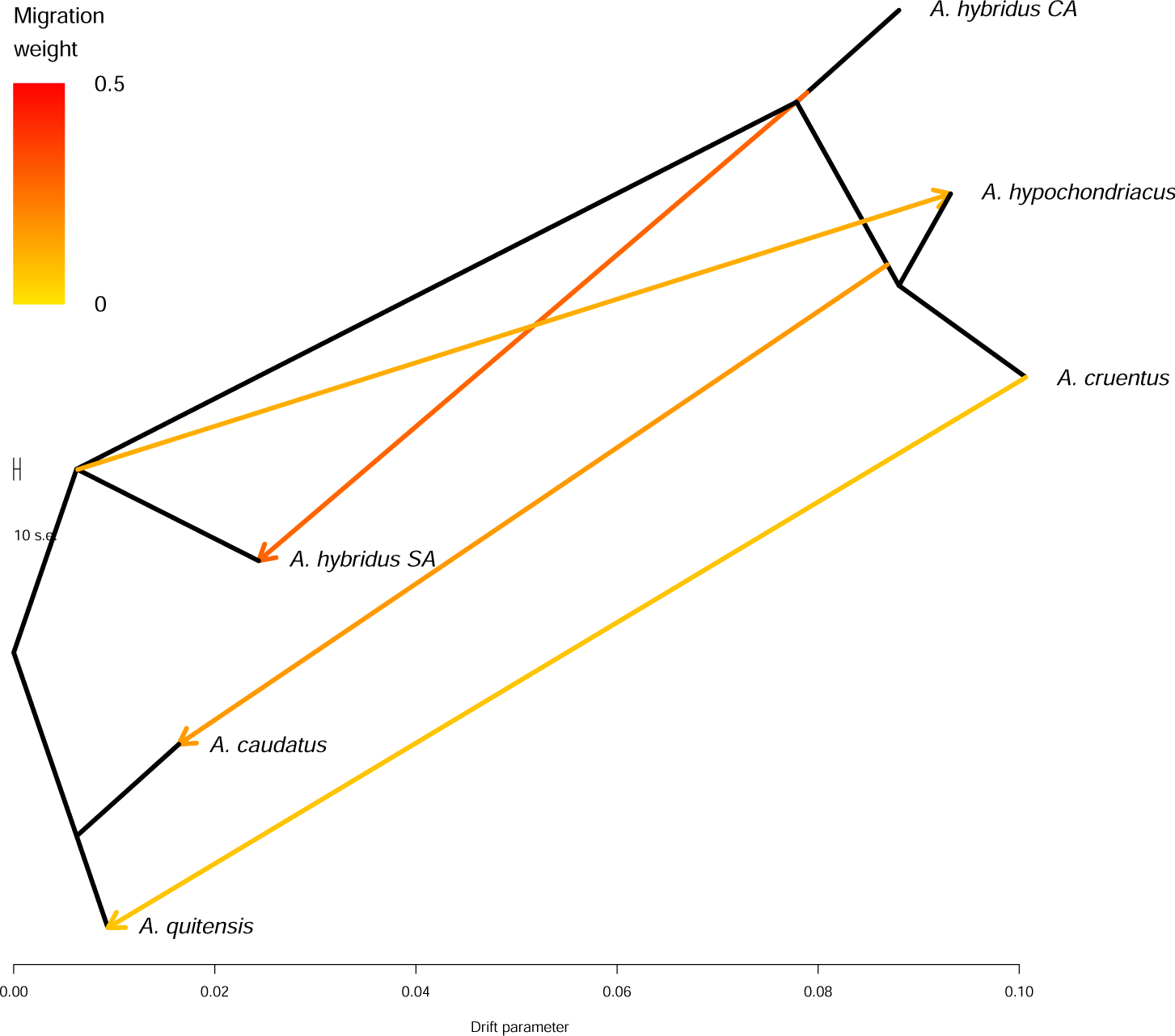
Allele frequency based species tree of wild (hybridus and quitensis) and domesticated amaranth (caudatus, cruentus and hypochondriacus). Arrows indicate gene flow between groups and their color indicates the intensity.

Despite the signals of gene flow between species, our results suggest that each crop species was selected from hybridus separately. In contrast to other crops, for which a single initial domestication processes followed by gene flow is supported (Choi and Purugganan 2018), we did not find shared signals of selection. For instance, two domestications had been the prevalent model for Asian rice (Huang *et al*. 2012), but recent analyses favor a single domestication followed by gene flow that resulted in a second crop subspecies (Choi and Purugganan 2018). While we find similar values of genome-wide differentiation (F_*ST*_) between wild and domesticated populations as reported for Asian rice (Huang *et al*. 2012), there was no overlap of selective sweeps between the grain amaranths (Figure 3D) as observed in Asian rice (Choi and Purugganan 2018). From our results we cannot exclude the possibility that other domestication related genes were contributed through gene flow between populations, but haplotypes of the seed color QTL showed a distinct pattern for each grain species that was closer to a group of wild amaranth than to other grain species (Figure S10 and S11), suggesting an independent selection for this trait rather than initial selection followed by gene flow.

The demographic history of crop species influences the extent and type of phenotypic variation available for selection during domestication. Polygenic domestication traits like grain size or seed shattering may require evolutionary changes in multiple loci (Huang *et al*. 2010; Xue *et al*. 2016) and more genetic variation to select the favorable combination for a functional domestication trait. A potential reason for incomplete domestication in amaranth might be a lack of functional diversity for complex traits due to a small population size before or during domestication. We therefore reconstructed the demographic history and the split times for the different species.

The demographic analysis showed that all five species followed similar trajectories before the split, with minor differences in the timing of demographic changes (Figure S18). Hybridus had a larger effective population size (N_e_) than the three domesticated species, which was consistent with the level of diversity of present populations (Figure 1A and B). Estimated split times of domesticated amaranths from their wild ancestor hybridus ranged from 50,000 generations for caudatus to 8,500 generations in hypochondriacus. In South America, the wild relative quitensis split more recently from hybridus (35,000 generations) than the domesticated caudatus. The decline in population size in hybridus predated the split of the species and propagated into all species (Figure 5 and S18).

**Figure 5.**
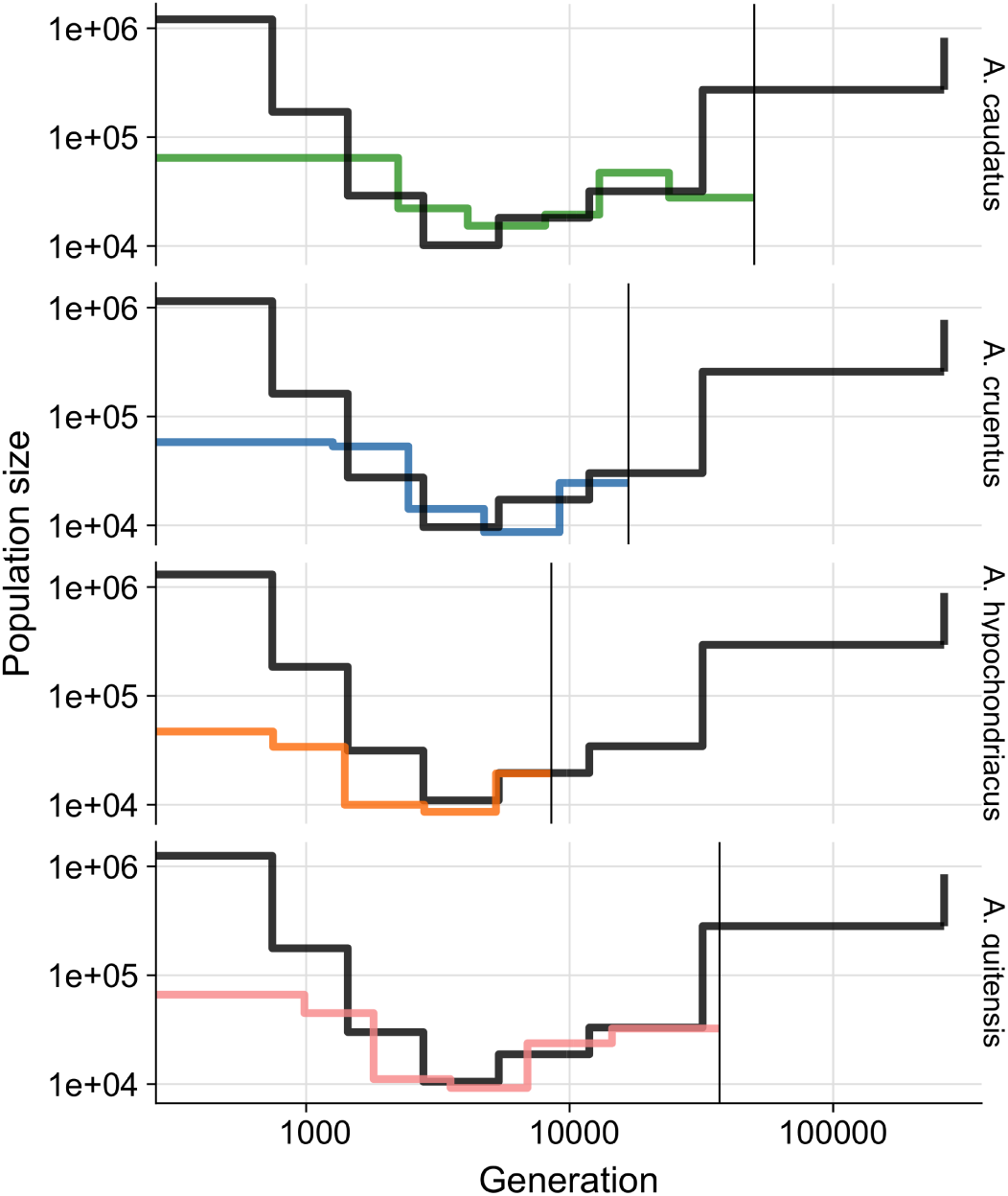
Demographic history of grain amaranth species. Black lines show population size estimates for hybridus. Vertical line shows the estimated split time between wild ancestor and domesticated amaranth in generations before present.

The similarity of population histories before domestication-related splits between species indicated high confidence for the demographic model. The early date estimated as split time for caudatus might have been be confounded by introgression from Central American amaranth into caudatus (Figure 4) and a potentially inaccurate estimate of the mutation rate, which was taken from *A. thaliana*. While the mutation rate influences the absolute datings, it does not influence the relative timing between populations. The independent demographic analyses suggest that the bottleneck in amaranth predates its domestication, leading to reduced population size in the ancestor of the three crop species during the time of domestication (Figure 5).

A bottleneck may have caused a lack of functional diversity for complex domestication traits during the time of domestication. Polygenic domestication traits, such as seed shattering and seed size, require either the presence of standing variation at functional sites or a large number of new mutations during domestication. While new mutations can lead to rapid adaptation for traits with major regulator genes, such as seed color, which we show here to be controlled by two major QTL (Figure 2), they are less likely to drive polygenic adaptation with a large number of small effect alleles (Stetter *et al*. 2018; Sella and Barton 2019). Hence, a small population size in the ancestor of all three grain amaranths that we observe here could potentially have prevented the adaptation of complex traits and might have interfered with a complete domestication in amaranth.

## Conclusion

In this study we revealed that amaranth was domesticated three times from a single ancestor (hybridus), twice in Central America and once in South America. Yet each of these attempts was incomplete, because key domestication traits like larger seeds are missing in all three grain amaranth species. We found phenotypic convergence for seed color conversion and identified the transcription factor MYBl1 as potential candidate gene for the seed coat color determination in amaranth. The seed color QTL regions did not show the signature of hard selective sweeps in the three domesticated amaranth species, similar to several domestication genes in other crops (Studer *et al*. 2011; Zhu *et al*. 2012). However, in hypochondriacus we identified signals of a soft selective sweep within the QTL region that contains AmMYBl1 (Figure S16). The white seed color trait in amaranth might have been selected from standing genetic variation in the wild ancestor, where the recessive white alleles are probably deleterious. In other crop ancestors dark seeds provide resistance to biotic stresses, but are selected against during domestication, because of human color preferences and pleiotropic effects with other domestication traits (Sweeney and McCouch 2007).

Hundreds of crops show a similar lack of complex domestication traits as amaranth. Our case study of an incomplete domestication indicates that the population size decline in the ancestor might have resulted in a lack of diversity to simultaneously select several quantitative domestication traits even under strong artificial selection by humans. The demographic history has been reconstructed for a number of crops, which show bottlenecks during domestication, but have larger ancestral effective population size in the wild ancestors (Zhou *et al*. 2017; Beissinger *et al*. 2016; Wang *et al*. 2017; Cubry *et al*. 2018; Meyer *et al*. 2016). To further investigate this potential reason for incomplete domestication the demographic history of more incomplete domesticates should be studied and more potential reasons for a lack of adaptation, e.g., trait pleiotropy (Caspari 1952) and the balance between new beneficial mutations and genetic load (Morran *et al*. 2009) should be discovered. Crops and their domestication are an ideal model to study polygenic adaptation and understand how selection acts on multiple complex traits simultaneously and the genomic signal of allele frequency changes (Purugganan 2019). Understanding the domestication process, not only of major crops, but also the hundreds of locally grown crops is essential to adapt cropping systems and increase crop diversity for a sustainable future agriculture.

## Acknowledgments

We thank Julie Jacquemin and the Ross-Ibarra lab for helpful discussions, Elisabeth Kokai-Kota for technical assistance in DNA sequencing and David Brenner, Fabian Freund, Jeffrey Ross-Ibarra, Katharina Böndel, Michelle Stitzer and Philipp Brand for comments on earlier versions of the manuscript. We thank David Brenner and the USDA amaranth germplasm collection for the permission to use amaranth images in figure 1.

## Author Contributions

M.G.S and K.J.S designed the experiments and supervised the research. M.G.S produced the mapping population, M.G.S conducted population genetic analyses, M.V. and M.G.S conducted bulk segregant and QTL mapping analysis. M.G.S wrote the draft of the manuscript. All authors edited, read and agreed to the final version of the manuscript.

## Data Availability

Sequencing data is available from the European Nucleotide Archive (ENA) under the project numbers PRJEB30531 (whole genome diversity panel), PRJEB30532 (BSA data), and PRJEB30533 (F2 mapping population). Scripts used for the analysis are available through figshare (10.6084/m9.figshare.7600166)

## Methods

### Plant material and sequencing

Geographically diverse amaranth accessions (Table S1) representing all three grain amaranth species, *A. caudatus* L., *A. cruentus* L. and *A. hypochondriacus* L., and three wild relatives, *A. hybridus* L., *A. quitensis* L. and *A. powellii* S. Watson, were obtained from USDA ARS genebank. DNA of young leaves of a single plant from each accession was extracted using the DNeasy Plant Mini Kit (Qiagen, Netherlands) following manufacturers’ instructions and sequencing libraries were prepared after quality control. Samples were sequences on an Illumina NextSeq with 2×150 bp with VIB Genomics Core Facility (Belgium) or an HiSeq4000 with 2×100 bp by Macrogen (Korea).

### Data processing and filtering

Sequencing reads were filtered and trimmed with `Trimmomatic v0.36` (Bolger *et al*. 2014) and read quality was assessed with `fastQC` (https://www.bioinformatics.babraham.ac.uk/projects/fastqc/). High quality reads were mapped to the *A. hypochondriacus* reference genome V2.1 (Lightfoot *et al*. 2017) using `BWA mem` (Li 2013). We compared the proportion of mapped reads using an ANOVA and did not find significant differences in mapped reads between species (Table S2. SNPs were called and filtered according to GATK “Best Practices” (McKenna *et al*. 2010). Additional filtering for a maximum of 30 % missing values at a site, and only SNPs from the 16 largest scaffolds of the reference genome (corresponding to the 16 chromosomes) was performed with `vcftools 0.1.14` (Danecek *et al*. 2011). For the GWAS analysis SNPs were filtered for a minor allele frequency (MAF) *>*0.05.

### Population genetic analysis

We used filtered SNP data for all population genetic analyses. Population structure was inferred with `ADMIXTURE` (Alexander *et al*. 2009) with cross validation to determine cluster numbers most strongly supported by the data. Principle component analysis was performed after LD filtering in 10,000 bp windows with an LD threshold of 0.3 using SNPrelate (Zheng *et al*. 2012). The population network was assessed with CONE (Community Oriented Network Estimation)(Kuismin *et al*. 2017). First, sites with missing data were removed as well as sites with MAF < 0.05. To select the optimal value of the tuning parameter we used the StARS model selection with 20 sampling steps of 2,634 SNPs each and 40 lambda parameters ranging from 0.5 to 0.005. We then used the complete filtered SNP set for the neighborhood selection procedure. The population network was constructed with the Fruchterman-Reingold algorithm and the additional StARS weights were used to weight the edges. To test for robustness, we used the same procedure in 20 random SNP samples of 10,000 and 20,000 SNPs.

Nucleotide diversity (*π*) and Tajima’s *D* (Tajima 1989) were calculated using `vcftools 0.1.14` (Danecek *et al*. 2011). *F*_*ST*_ values were calculated with SNPrelate (Zheng *et al*. 2012).

### Mapping population and QTL mapping

An F_2_ mapping population was created from a single F_1_ hybrid plant between ‘Oscar Blanco’ (PI 642741,*A. caudatus*, white seeds) and PI 490705 (*A. quitensis*, dark seeds). A total of 224 F_2_ plants were genotyped using the original protocol for genotyping by sequencing (GBS) (Elshire *et al*. 2011) and their seed color was recorded at maturity after cultivation in a greenhouse. DNA of young leaves was extracted and GBS libraries were prepared as described in (Stetter *et al*. 2017b) and sequenced on a Illumina HiSeq4000 instrument with 2×100 bp by Macrogen. After read filtering, alignment the *A. hypochondriacus* reference genome V2.1 (Lightfoot *et al*. 2017) with `BWA mem` (Li 2013) and SNP calling with `samtools` (Li *et al*. 2009) and 206 samples were kept for further analysis. SNP data was filtered to allow a maximum of 70% missing values at a site and minor allele frequency ≤ 0.01. Linkage mapping was performed with R/qtl (Arends *et al*. 2010).

### Bulk segregant analysis

Leaf samples from a total of 200 plants with contrasting seed colors (dark and white) were collected from the Hohenheim Amaranth Breeding Population in October 2016 in Stuttgart, Germany. This advanced population is an early breeding population (three mass selection cycles) consisting of segregating populations from spontaneous (interspecific) hybrids and genetic resources mainly consisting of *A. cruentus* and *A. hypochondriacus*. One hundred single plants for each seed color were collected from diverse parental backgrounds to form two bulks. DNA was extracted using the DNeasy Plant Mini Kit (Qiagen, Netherlands) following manufacturers instructions and TruSeq DNA PCR-Free Library (Illumina, USA) sequencing libraries were prepared after quality control. Libraries were sequenced with 2×150 bp on a Illumina HiSeq X instrument by Macrogen at 23-28x coverage. After quality control with `FastQC`, reads were trimmed with `Trimmomatic v0.36` (Bolger *et al*. 2014) and mapped to the *A. hypochondriacus* reference genome V2.1 (Lightfoot *et al*. 2017) with `BWA mem` (Li 2013) and SNPs were called according to GATK “Best Practices” (McKenna *et al*. 2010). Bulk segregant analysis was performed with the G’ method (Magwene *et al*. 2011) implemented in QTLseqr R library Mansfeld and Grumet (2018) on a total of 2,656,505 sites after filtering for high quality SNPs.

### Genome wide association mapping and candidate gene identification

Seed colors were inferred visually from seed samples for the 121 diverse individuals and coded binary. We performed GWAS on 276,466 LD pruned SNPs that were used for the PCA for 116 unambiguous white and dark seed accessions (accessions with mixed seed colors were removed from the analysis) using the compressed mixed linear (Zhang *et al*. 2010) model implemented in GAPIT (Lipka *et al*. 2012) and chose a cut-off p-value of 10^−5^. We compared the CDS regions of the hypochondriacus reference genome (Lightfoot *et al*. 2017) which was generated from an individual with white seeds to that of its distant wild relative *A. tuberculatus* (Kreiner *et al*. 2019) with dark seeds. We used the CoGe synmap online tool (genomevolution.org/coge/SynMap.pl) to align CDS regions and downloaded the results for plotting and annotating genes in the QTL region. To estimate linkage disequilibrium within the QTL region in each speices and color group, we calculated pairwise R^2^ values between sites using `–ld` (Gaunt *et al*. 2007) implemented in PLINK 1.9 (www.cog-genomics.org/plink/1.9/) for each species and seed color group. To identify independent or shared origin of the seed color QTLs we performed a PCA analysis on SNPs within the two QTL regions on chromosome 3 and 9 identified using mapping and GWAS. Gene annotations and sequences were extracted from the *A. hypochondriacus* reference genome V2.1 (Lightfoot *et al*. 2017). The impact of variants within AmMYBl1 region was annotated and summarized using snpEff (Cingolani *et al*. 2012). Haplotypes were plotted for the region around AmMYBl1 (chr 9: 1405000 – 14709000) and the entire QTL region on chromosome 3 (chr 3: 5881516 – 5966304) using haplostrips (Marnetto and Huerta-Sánchez 2017).

Additionally we performed a reference-free association study of *k*-mers using HAWK (Hitting Associations With *K*-mers) (Rahman *et al*. 2018). We ran the pipeline for each of the three main clusters identified in the population network of CONE separately using the seed color (excluding accessions with ambiguous color) as binary trait (case-control), and a *k*-mer length of 32bp. As part of the HAWK pipeline, significantly associated *k*-mers were assembled into contigs using ABySS (Simpson *et al*. 2009). Contigs were then mapped to the amaranth reference genome to determine their genomic position.

### Selective sweep detection

We identified regions with a signature of selective sweeps with a composite likelihood ratio test (Nielsen *et al*. 2005) implemented in SweeD (Pavlidis *et al*. 2013) by moving in 20 kb windows along the genome. We applied the test to each amaranth population and considered the top 5% of SNPs as outliers. To analyze convergence between the populations we compared the overlapping outliers between populations using upsetR (Lex *et al*. 2014) and tested if overlaps were significant through a 1,000,000 times repeated permutation unsing the sample.int function in R (R Core Team 2019). In addition, we analyzed each population using RAiSD (Alachiotis and Pavlidis 2018) with default parameters. To detect potential soft selective sweeps we calculated G12, G2/G1 and G123 statistics (Harris *et al*. 2018) which is able to detect hard and soft sweeps. We performed the analysis using public python scripts (https://github.com/ngarud/SelectionHapStats) and the three G statistics were calculated for each population and chromosome separately using a window size of 400 SNPs and a step size of 50 SNPs. Identification of G12 peaks was done using the default method implemented in H12peakFinder.py and selected afterwards the top 10 G12 peaks. We used as an indicator of a soft sweep a high value of both G2/G1 and G123 values on the top 10 G12 peaks and the comparison to the other two methods that detect hard selective sweeps.

### Gene flow and reconstruction of demographic history

We modeled the level of gene flow between amaranth populations with Treemix (Pickrell and Pritchard 2012) allowing for a maximum of four migration events. We compared residuals of the different models, which were strongly reduced with four migration events compared to zero migrations (Figure S17). To infer the demographic history of domesticated and wild amaranth populations we ran `SMC++` on the folded site frequency spectrum with a maximum gap size of 1,000 bp for each population separately and in pairs of *A. hybridus* and the respective other species to infer split times. As there is currently no reliably mutation rate estimate available for the different *Amaranthus* species we used the estimate for *Arabidopsis thaliana* of 7 × 10^−9^ (Ossowski *et al*. 2010). It should be noted that an incorrectly estimated mutation rate may offset the timescale of the population size inference and the estimated split time.

