## Supplementary figures and tables for "Parallel seed color adaptation during multiple domestication attempts of an ancient new world grain"

**Markus G Stetter**[*,†,1], **Mireia Vidal-Villarejo**[†] **and Karl J Schmid**[†,1]
[*]Botanical Institute, University of Cologne, Cologne, Germany, [†]Dept. of Plant Breeding, Population Genetics and Seed Science, University of Hohenheim, Stuttgart, Germany

**ABSTRACT**
Thousands of plants have been selected as crops, yet, only a few are fully domesticated. The lack of adaptation to agro-ecological environments of many crop plants with few characteristic domestication traits potentially has genetic causes. Here, we investigate the incomplete domestication of an ancient grain from the Americas, amaranth. Although three grain amaranth species have been cultivated as crop for millennia, all three lack key domestication traits. We sequenced 121 crop and wild individuals to investigate the genomic signature of repeated incomplete adaptation. Our analysis shows that grain amaranth has been domesticated three times from a single wild ancestor. One trait that has been selected during domestication in all three grain species is the seed color, which changed from dark seeds to white seeds. We were able to map the genetic control of the seed color adaptation to two genomic regions on chromosome 3 and 9, employing three independent mapping populations. Within the locus on chromosome 9, we identify a MYB-like transcription factor gene, a known regulator for seed color variation in other plant species. We identify a soft selective sweep in this genomic region in one of the crops species but not in the other two species. The demographic analysis of wild and domesticated amaranths revealed a population bottleneck predating the domestication of grain amaranth. Our results indicate that a reduced level of ancestral genetic variation did not prevent the selection of traits with a simple genetic architecture but may have limited the adaptation of complex domestication traits.

**KEYWORDS** Domestication, parallel evolution, orphan crop, MYB transcription factor, amaranth, crop wild relatives

### Supplement

#### Supplementary Figures

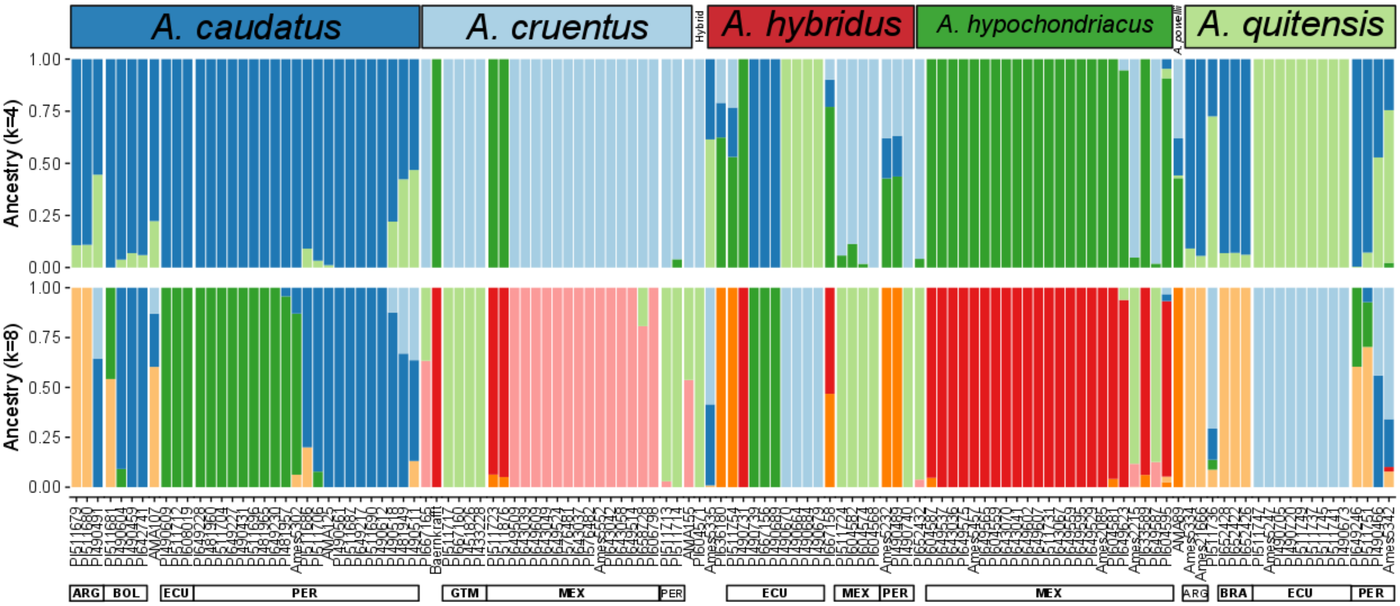


**Figure S1** Individual based ancestry contribution calculated with ADMIXTURE for $K = 4$, for three potential crop populations and wild ancestors and $K = 8$ representing species by geographic origin.

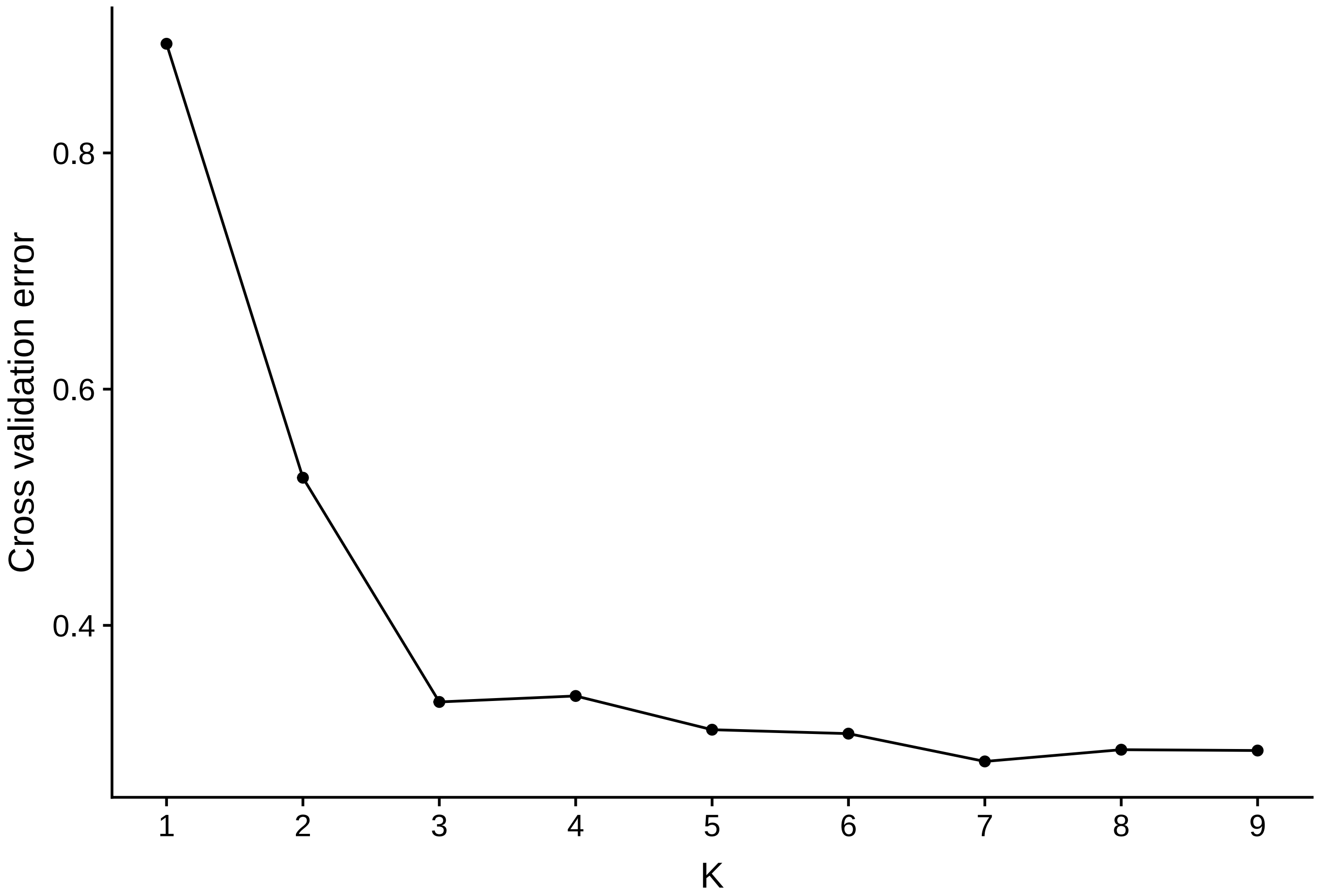


**Figure S2** Cross validation of ADMIXTURE.

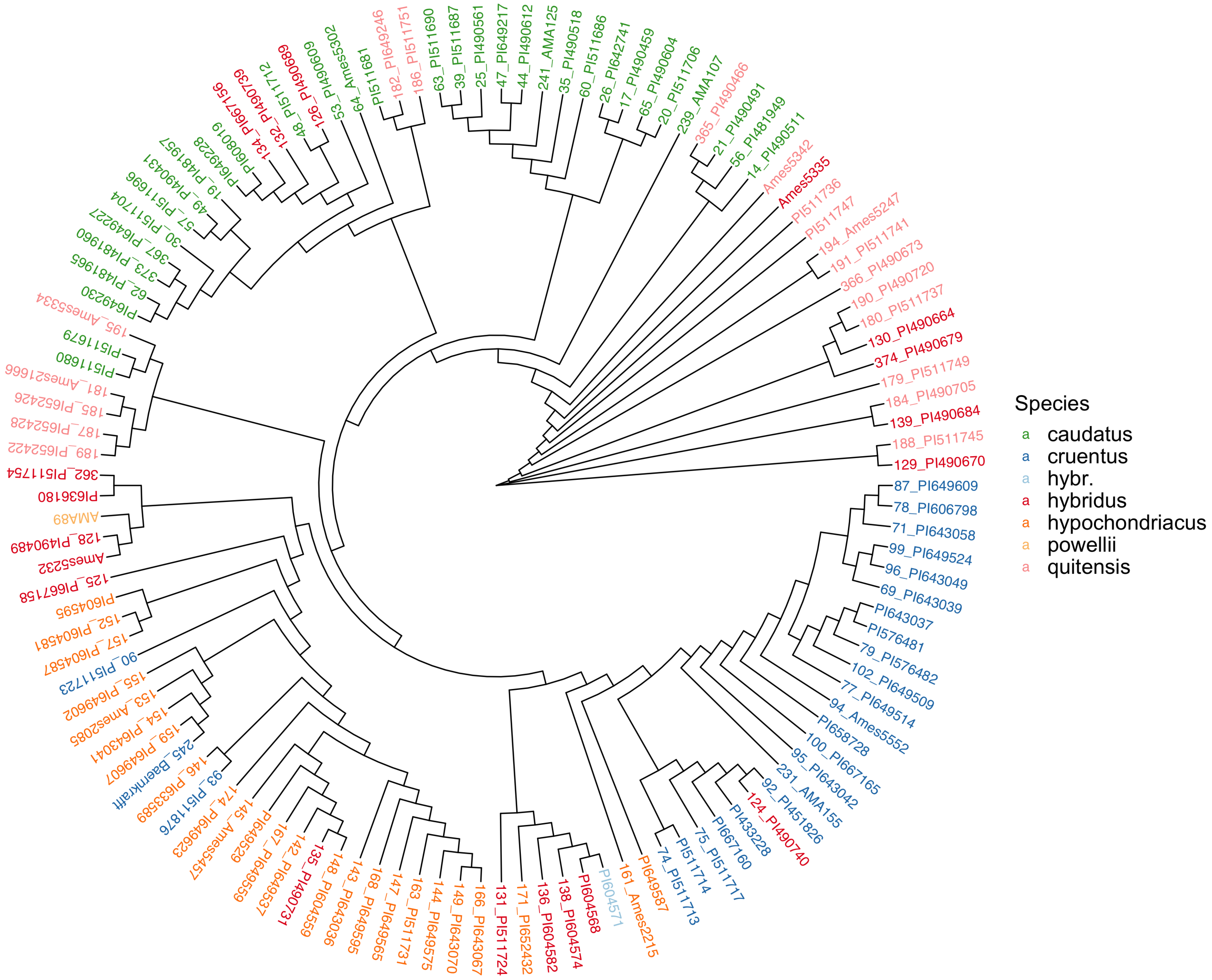


**Figure S3** Neighbor joining tree from genomic SNPs of 121 diverse individuals. Colors represent the different species.

**Figure S4 PCA** Interactive 3D representation of PCA showing PC1 - PC3.

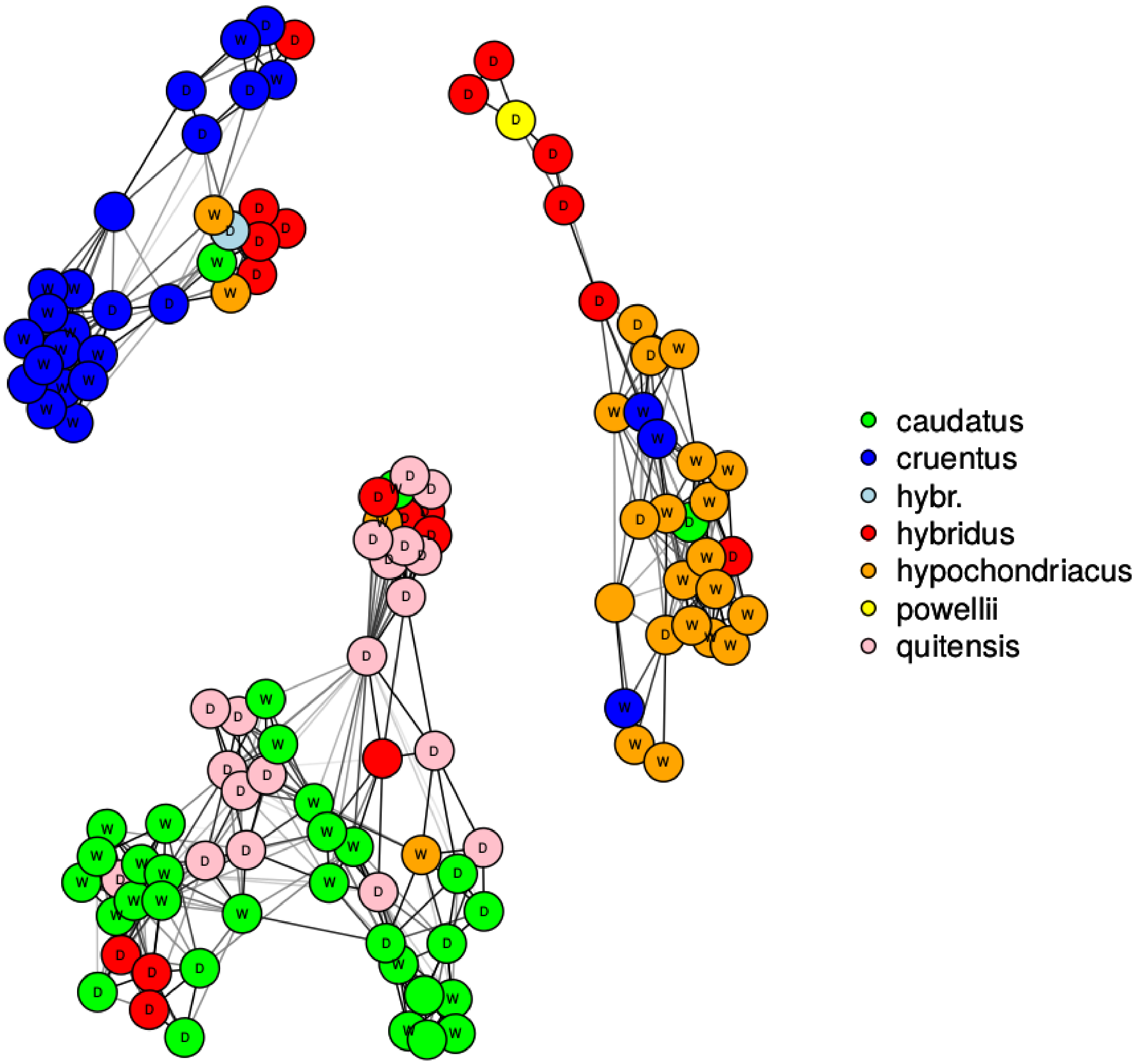


**Figure S5** Population network of CONE. Colors represent the different species. W indicates accessions with white seed color, D indicates dark seed color, nodes with no letter are accessions with ambiguous seed color.

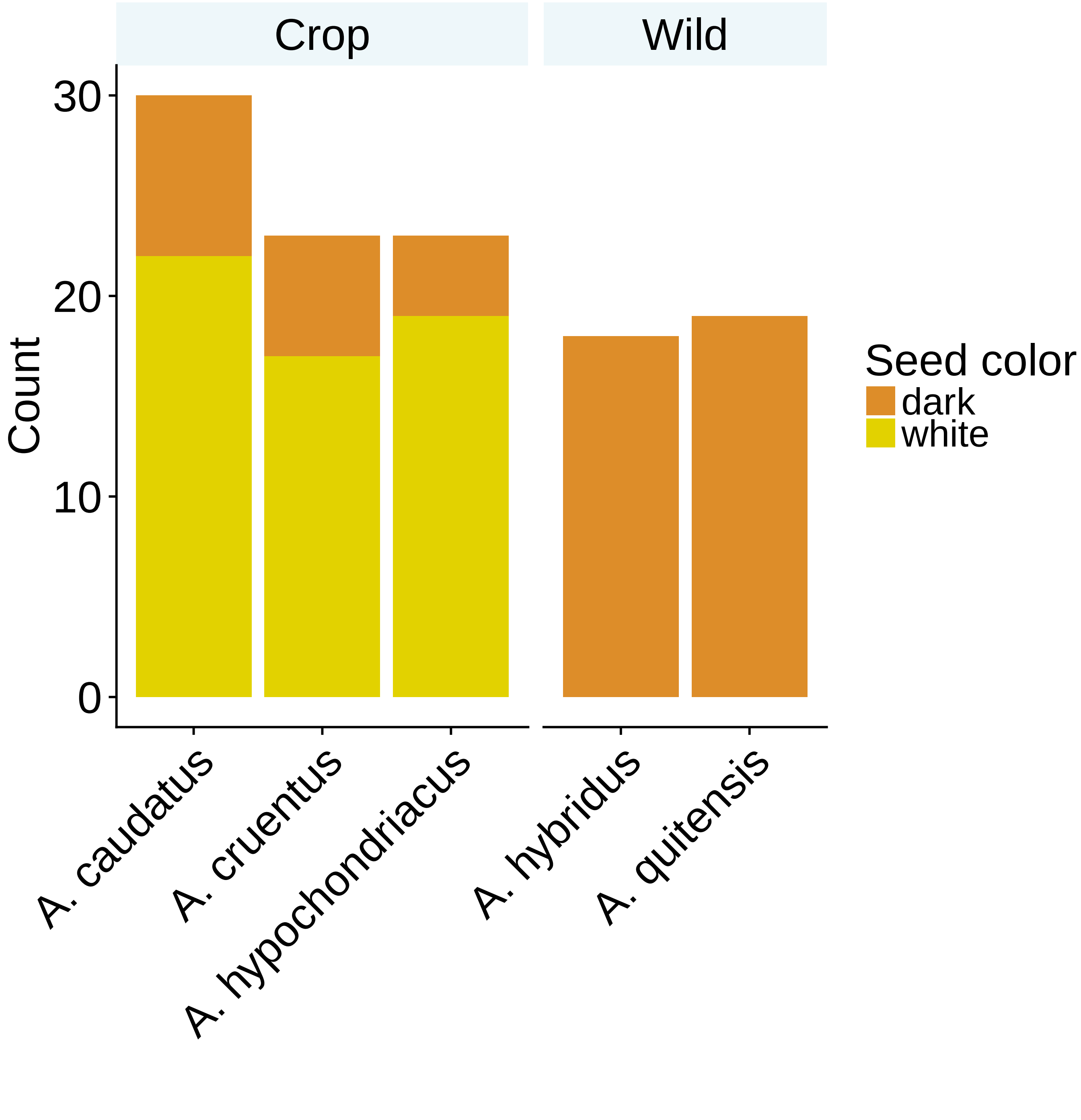


**Figure S6** Seed color phenotypes of diversity panel by species.

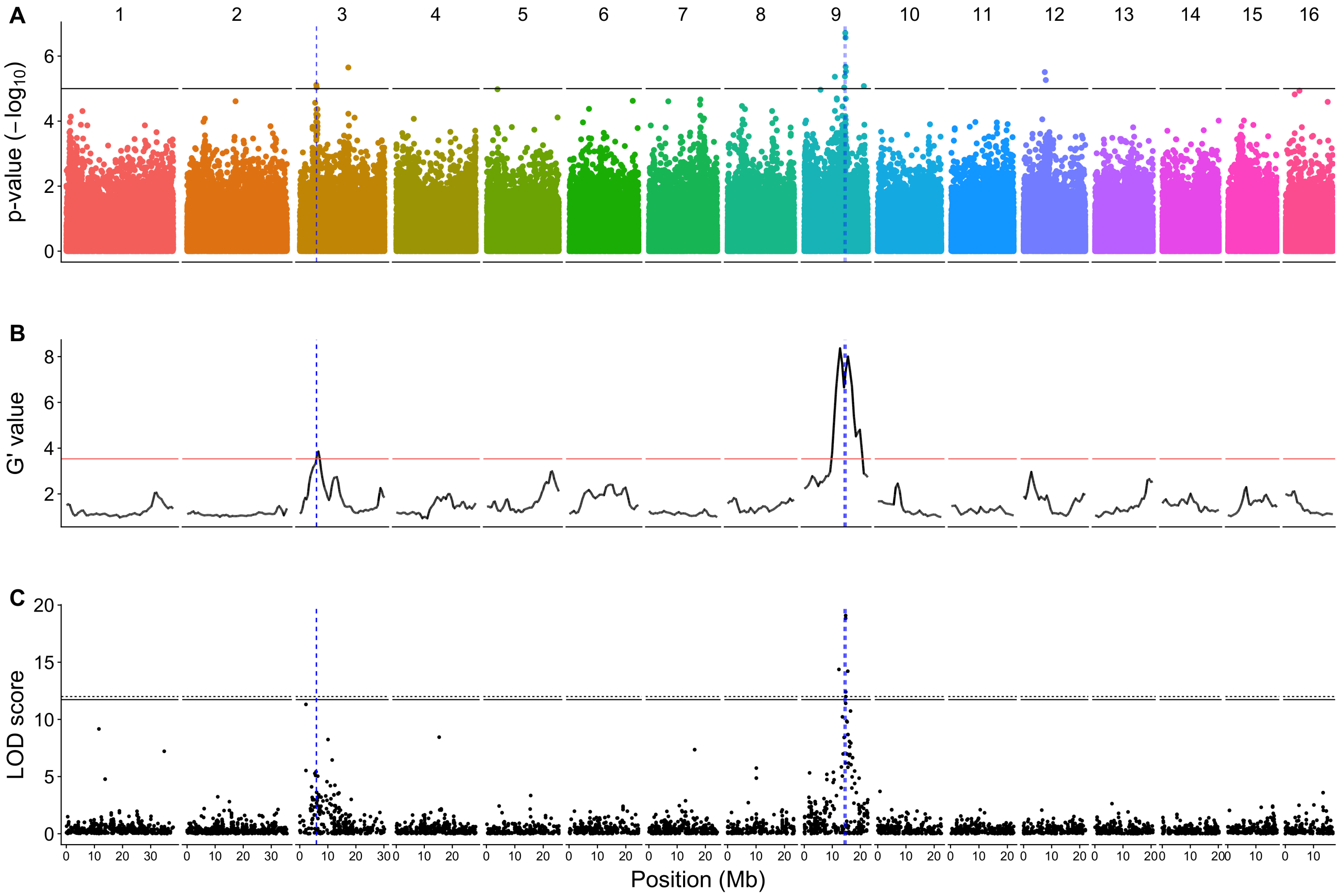


**Figure S7** Comparison of QTL regions across mapping populations. A) Genome wide association mapping in 116 diverse germplasm accessions, including crop and wild species B) Results of a bulked segregant analysis for seed color in 100 bulked samples of dark seeded individuals and 100 samples with white seeds from breeding population consisting of hypochondriacus and cruentus. C) QTL mapping results for seed color of $F_2$ mapping population between caudatus (white seeds) and quitensis (dark seeds).

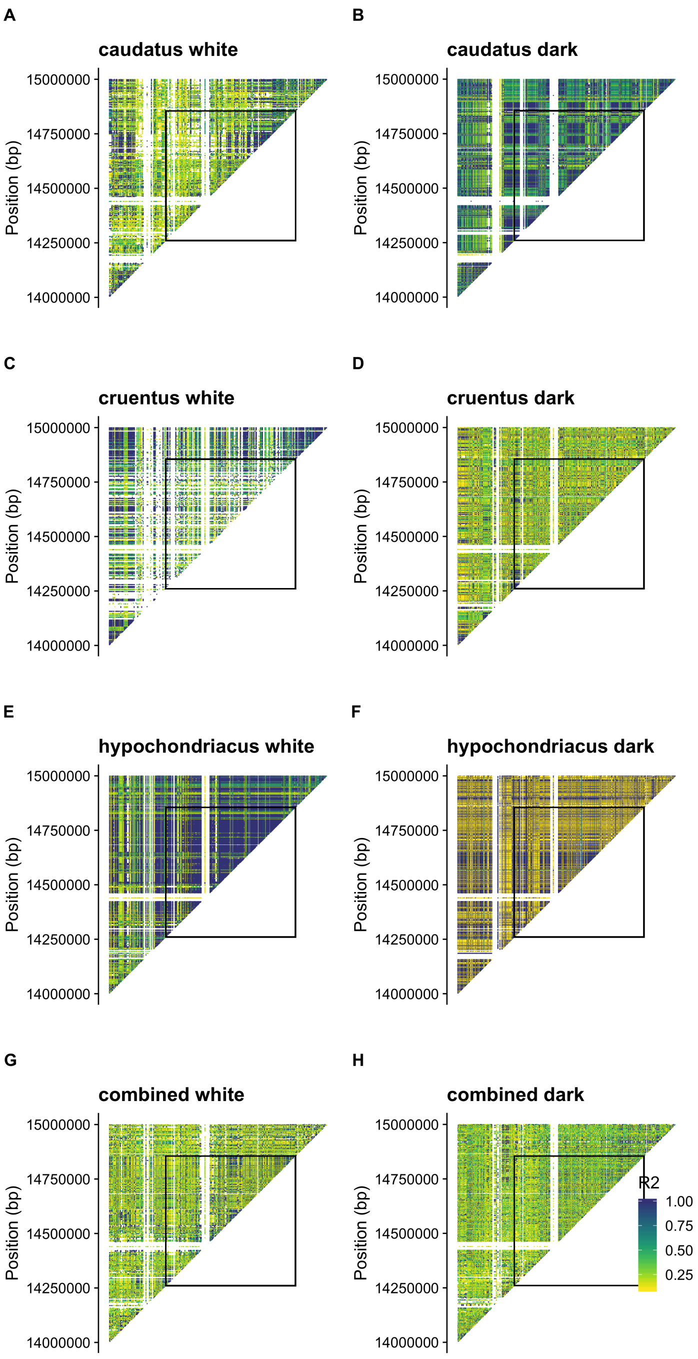


**Figure S8** Heat map of $R^2$ between sites within groups of dark and white seeds. Boxes indicate the location of the seed color QTL (table S3). Note that "hypochondriacus dark" includes only four samples (table S1)

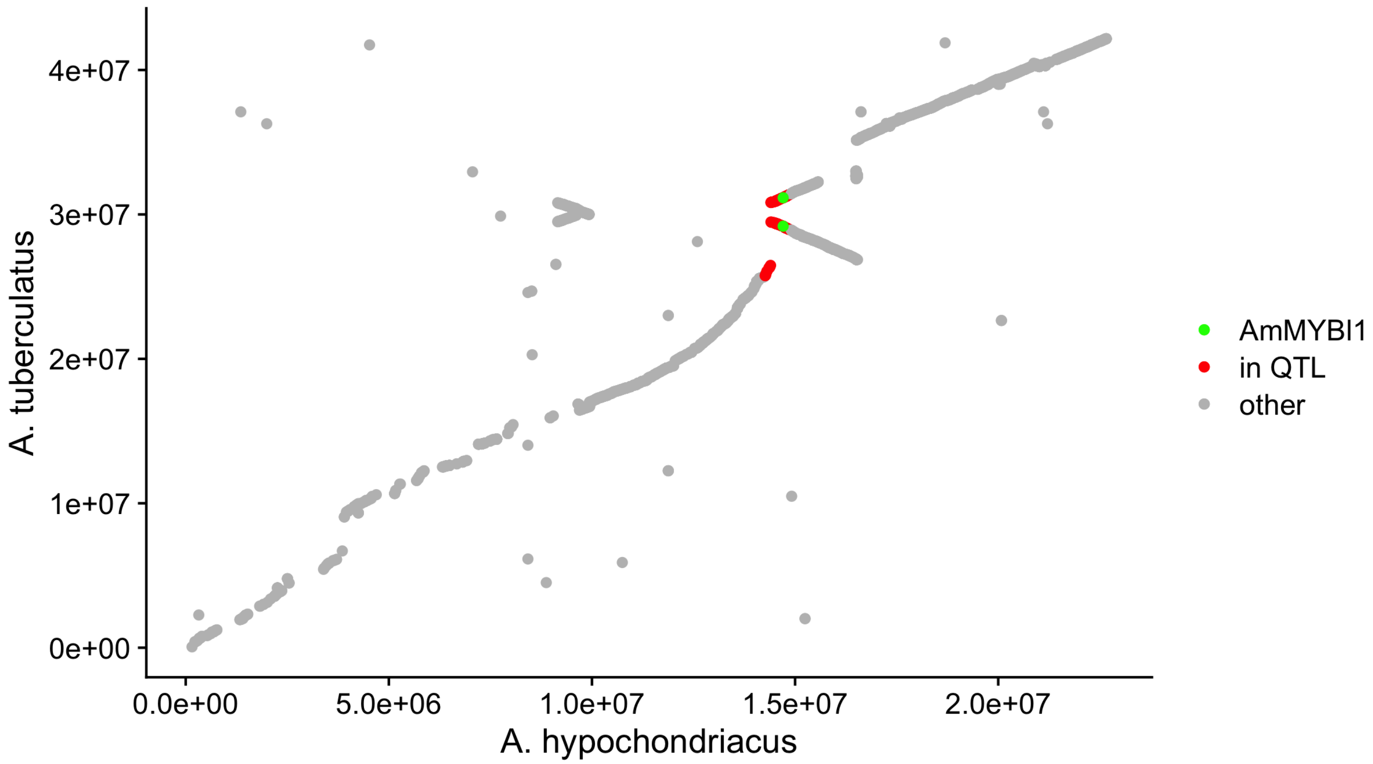


**Figure S9** Alignment of chromosome 9 of *A. hypochondriacus* and *A. tuberculatus* CDS regions. Indicating an inverted segment containing the seed color QTL (table S3)

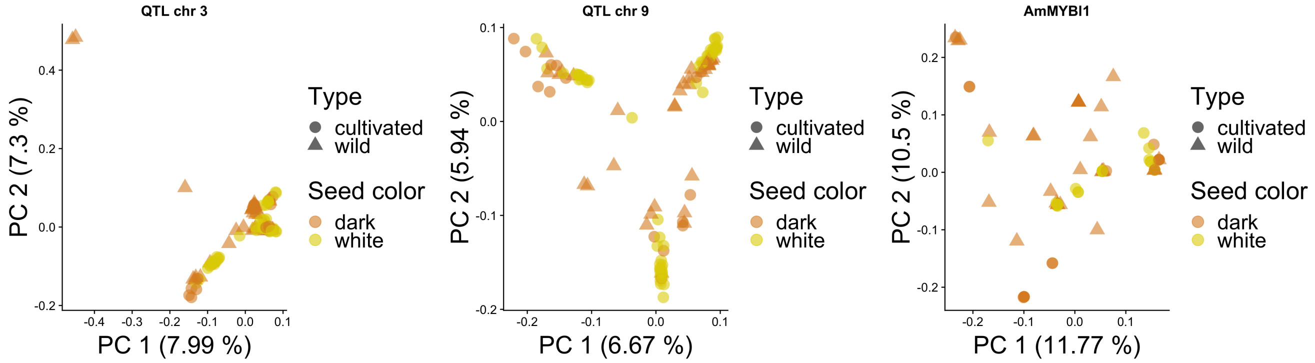


**Figure S10** Local PCA for each QTL region. A) QTL on chromosome 3 using 127SNPs after LD pruning, B) QTL on chromosome 9 using 933 SNPs after LD pruning in the region and C) AmMYBl1 region with 2 kb upsream and downstream

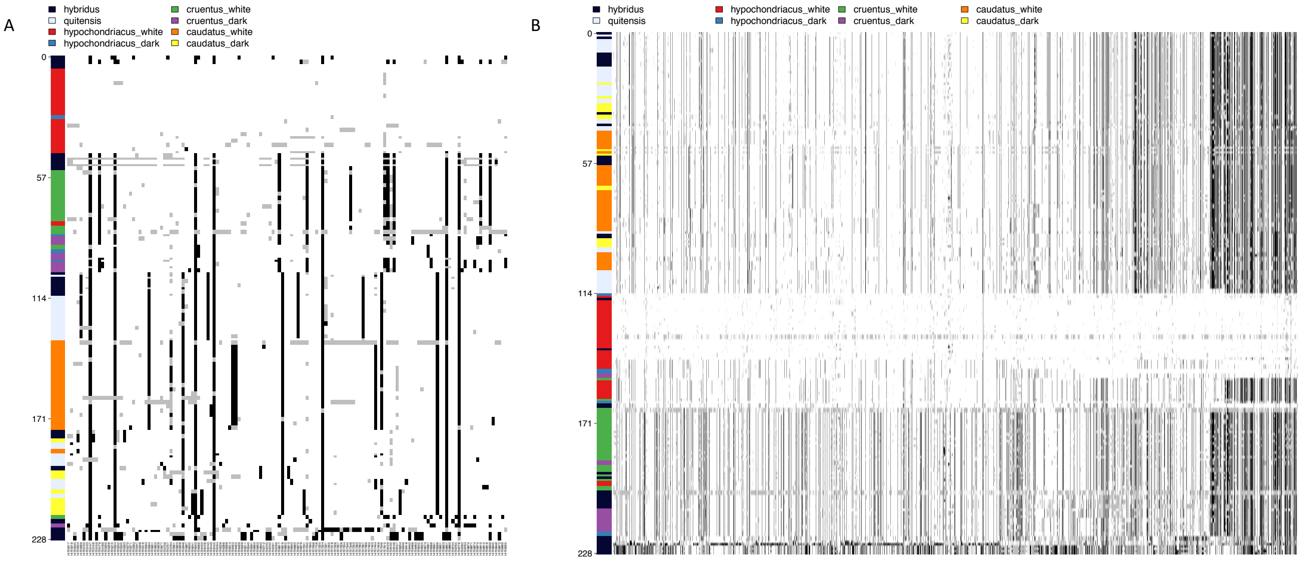


**Figure S11** Haplotype representation of SNPs in A) AmMYBl1 region with 2 kb upsream and downstream of the gene and B) QTL region on chromosome 3. White bars represent the reference allele, black bars the alternative allele and grey bars represent missing SNP values

**Number variants by type**

| Type | Total |
|---|---|
| SNP | 375 |
| MNP | 0 |
| INS | 53 |
| DEL | 54 |
| MIXED | 0 |
| INV | 0 |
| DUP | 0 |
| BND | 0 |
| INTERVAL | 0 |
| Total | 482 |

**Number of effects by impact**

| Type (alphabetical order) | Count | Percent |
|---|---|---|
| HIGH | 2 | 0.175% |
| LOW | 8 | 0.698% |
| MODERATE | 9 | 0.785% |
| MODIFIER | 1,127 | 98.342% |

**Number of effects by functional class**

| Type (alphabetical order) | Count | Percent |
|---|---|---|
| MISSENSE | 9 | 69.231% |
| SILENT | 4 | 30.769% |

Missense / Silent ratio: 2.25

**Number of effects by type and region**

Type

| Type (alphabetical order) | Count | Percent |
|---|---|---|
| downstream_gene_variant | 551 | 47.788% |
| frameshift_variant | 2 | 0.173% |
| intergenic_region | 322 | 27.927% |
| intron_variant | 145 | 12.576% |
| missense_variant | 9 | 0.781% |
| splice_region_variant | 6 | 0.52% |
| stop_gained | 1 | 0.087% |
| synonymous_variant | 4 | 0.347% |
| upstream_gene_variant | 113 | 9.801% |

Region

| Type (alphabetical order) | Count | Percent |
|---|---|---|
| DOWNSTREAM | 551 | 48.08% |
| EXON | 14 | 1.222% |
| INTERGENIC | 322 | 28.098% |
| INTRON | 141 | 12.304% |
| SPLICE_SITE_REGION | 5 | 0.436% |
| UPSTREAM | 113 | 9.86% |

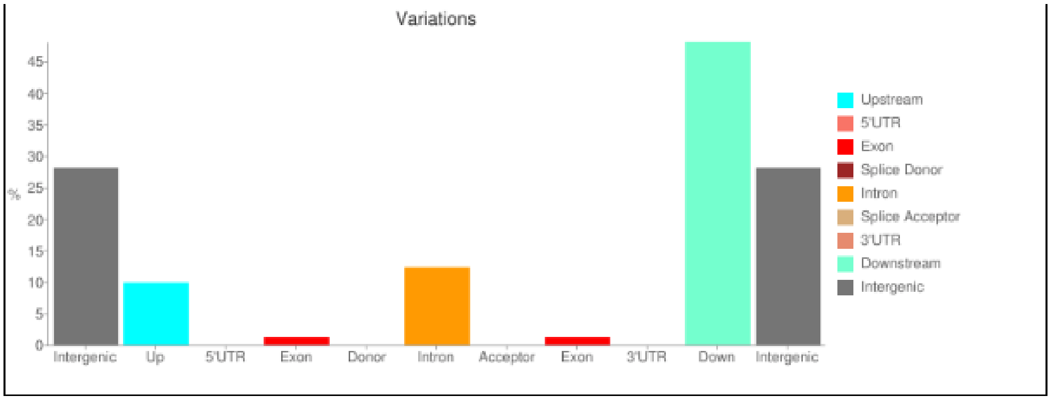


**Figure S12** Summary of variant impacts in AmMYBl1 including 2 kb up- and downstream of the gene.

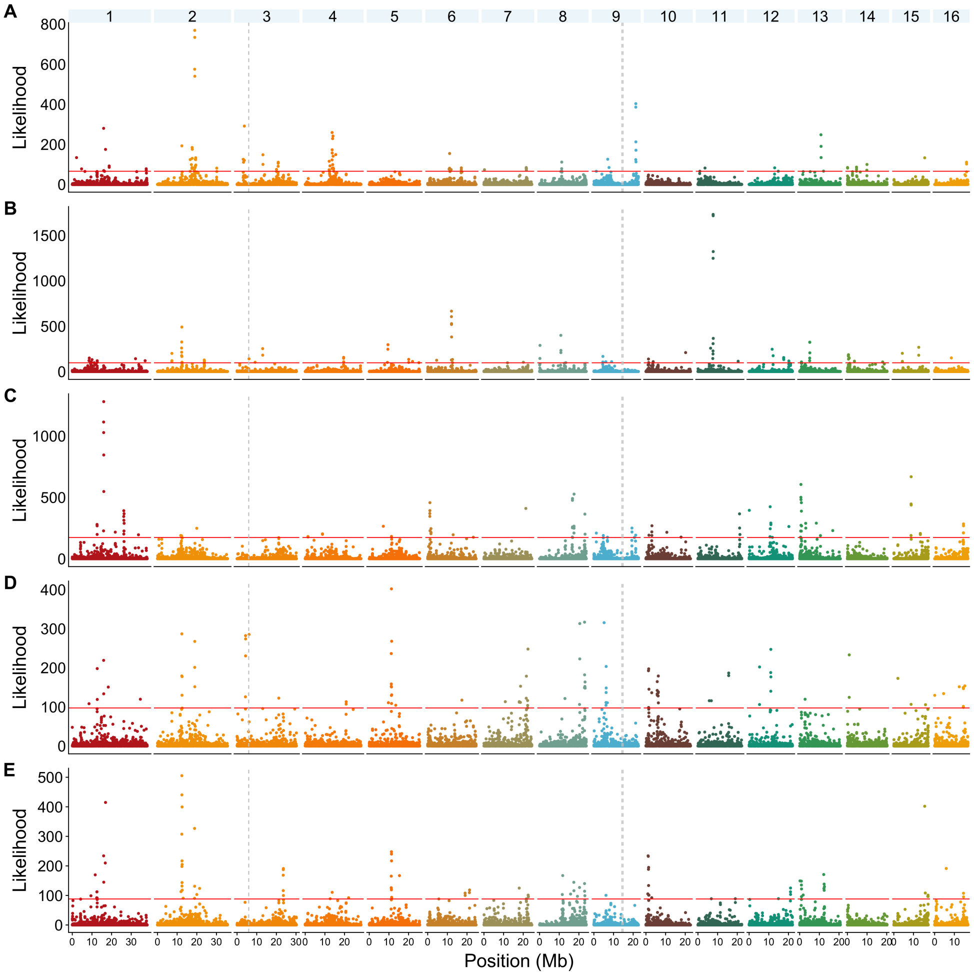


**Figure S13** Hard selective sweeps identified with SweeD in cultivated and wild amaranth populations along the genome. A) caudatus, B) cruentus, C) hypochondriacus, D) hybridus, E) quitensis. Horizontal line shows top 0.5% of cutoff for outliers

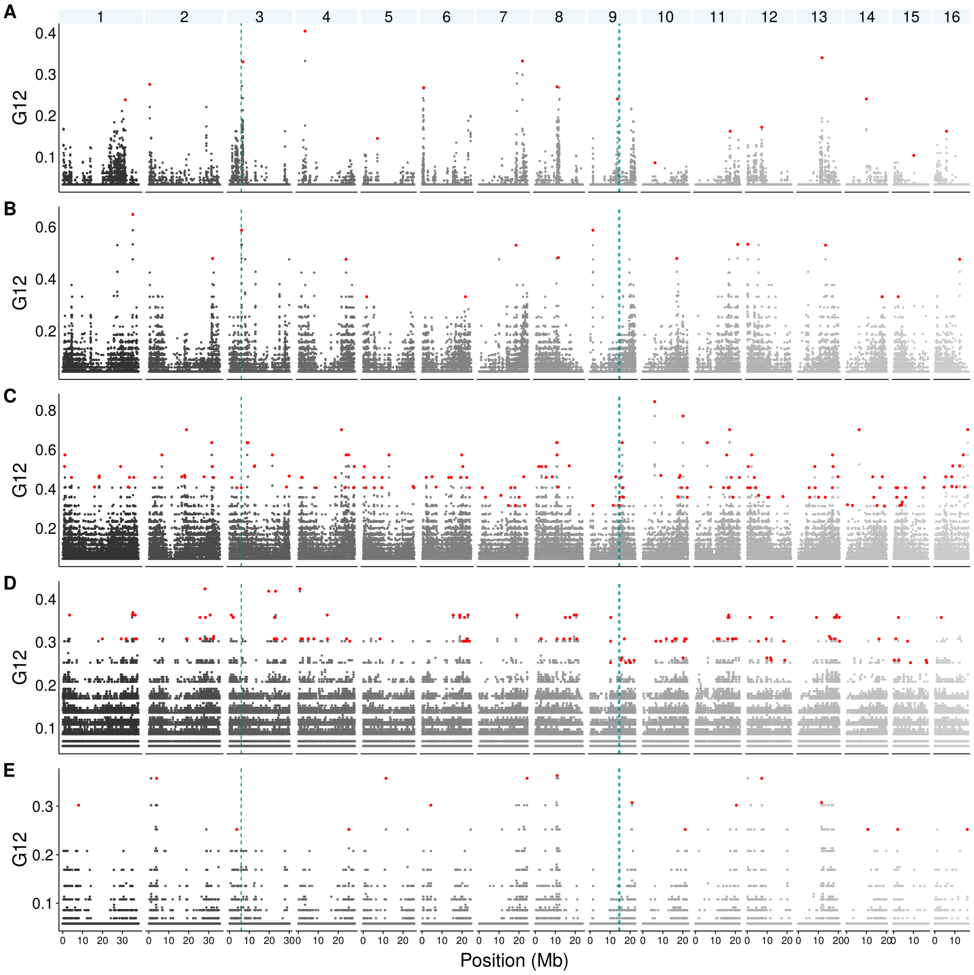


**Figure S14** Hard and soft selective sweeps as identified with the G2/G1 method in cultivated and wild amaranth populations along the genome. A) caudatus, B) cruentus, C) hypochondriacus, D) hybridus, E) quitensis. In red, highlighted the top 10 G12 peaks in each chromosome.

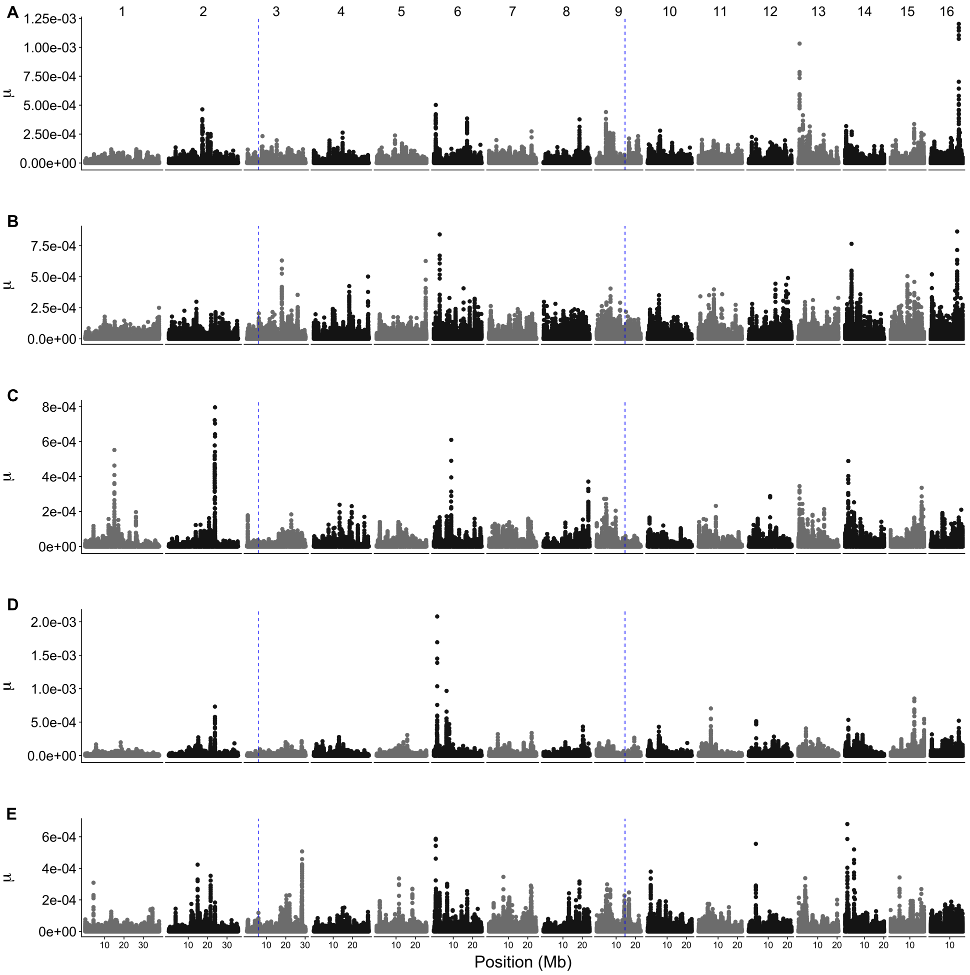


**Figure S15** Hard selective sweeps as identified with RAiSD method in cultivated and wild amaranth populations along the genome. A) caudatus, B) cruentus, C) hypochondriacus, D) hybridus, E) quitensis. Horizontal line shows top 0.5% of cutoff for outliers

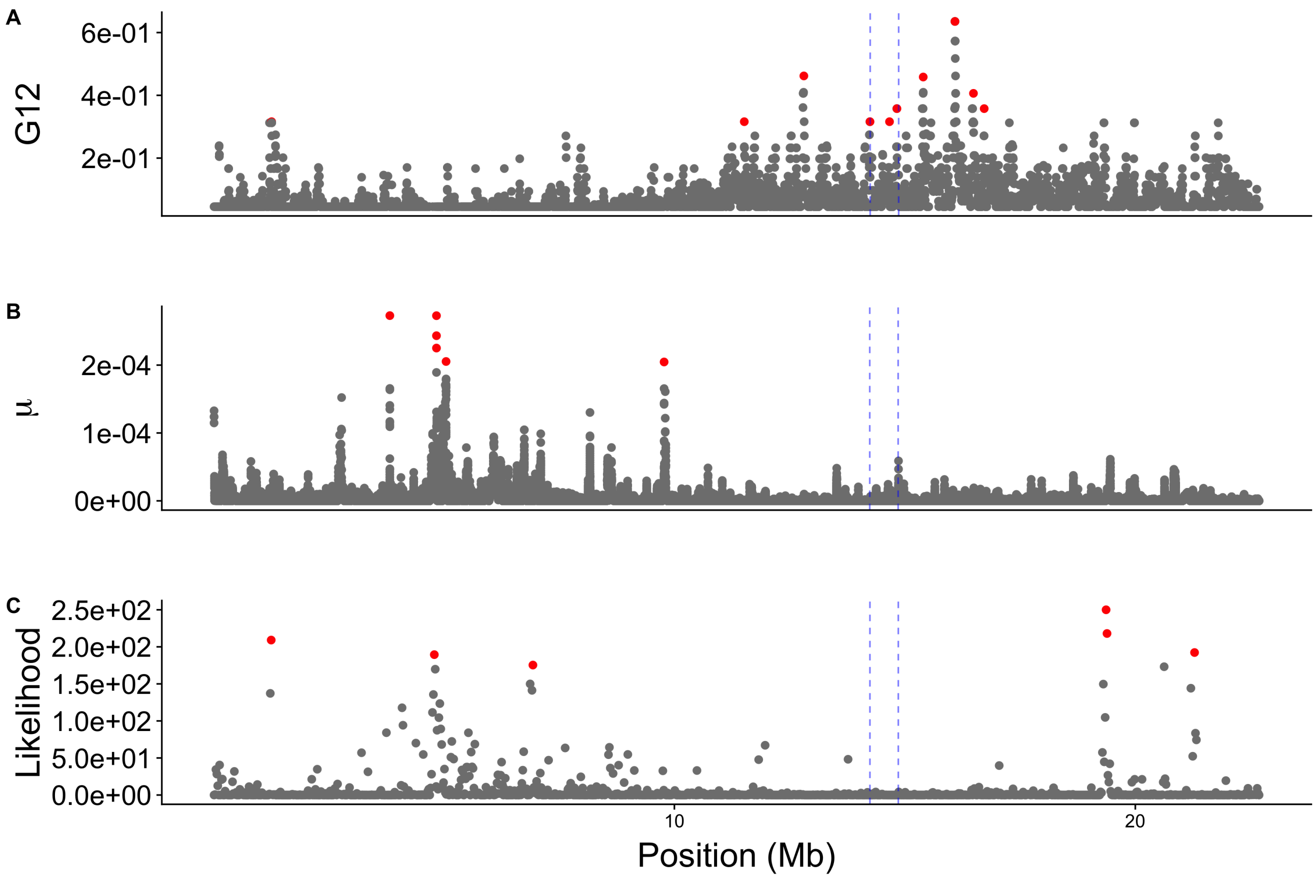


**Figure S16** Top 10 selective sweep hits in hypochondriacus on Chromosome 9 using three selection tests, A) G2/G1, B) RAiSD and C) SweeD. G2/G1 identifies hard and soft selective sweeps while the other two methods only identify hard sweep signals.

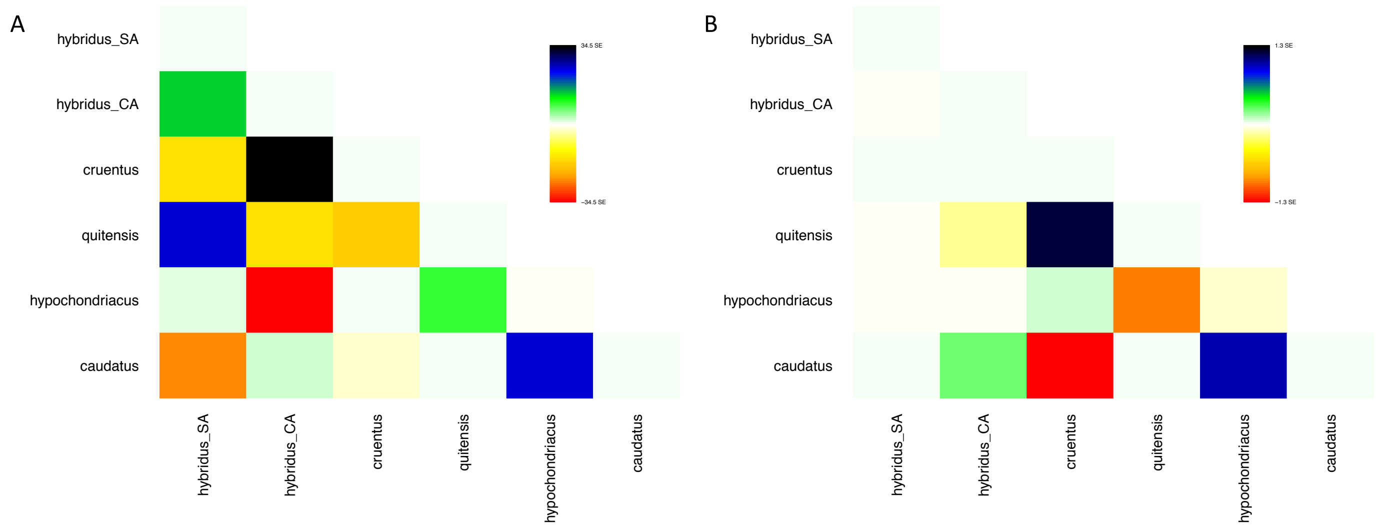


**Figure S17** Treemix residuals for trees with A) no migration event and B) with four migration events.

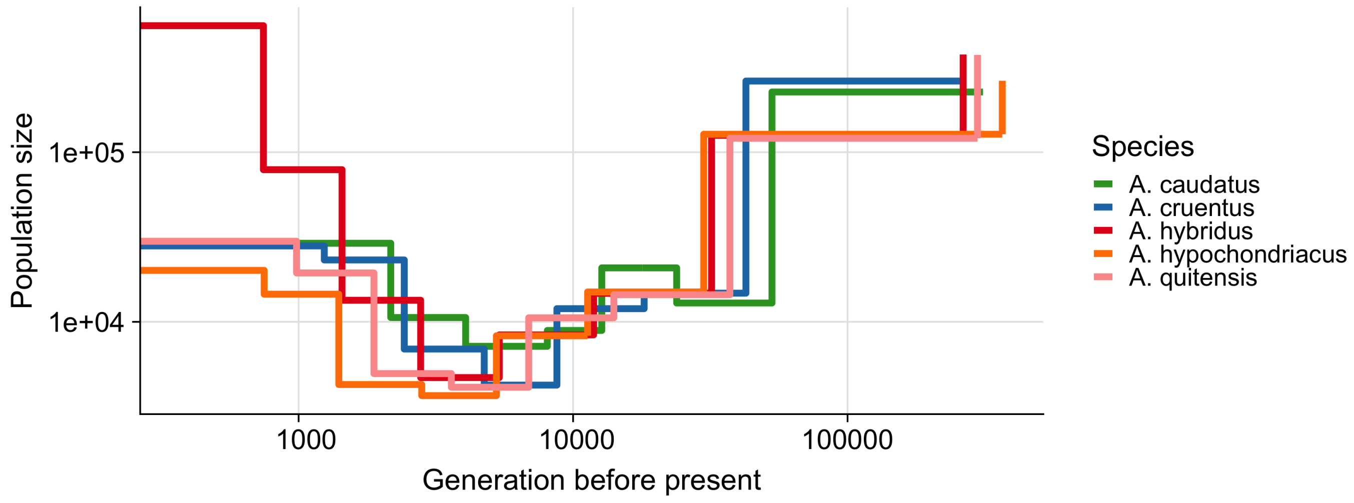


**Figure S18** Demographic history for individual amaranth populations.

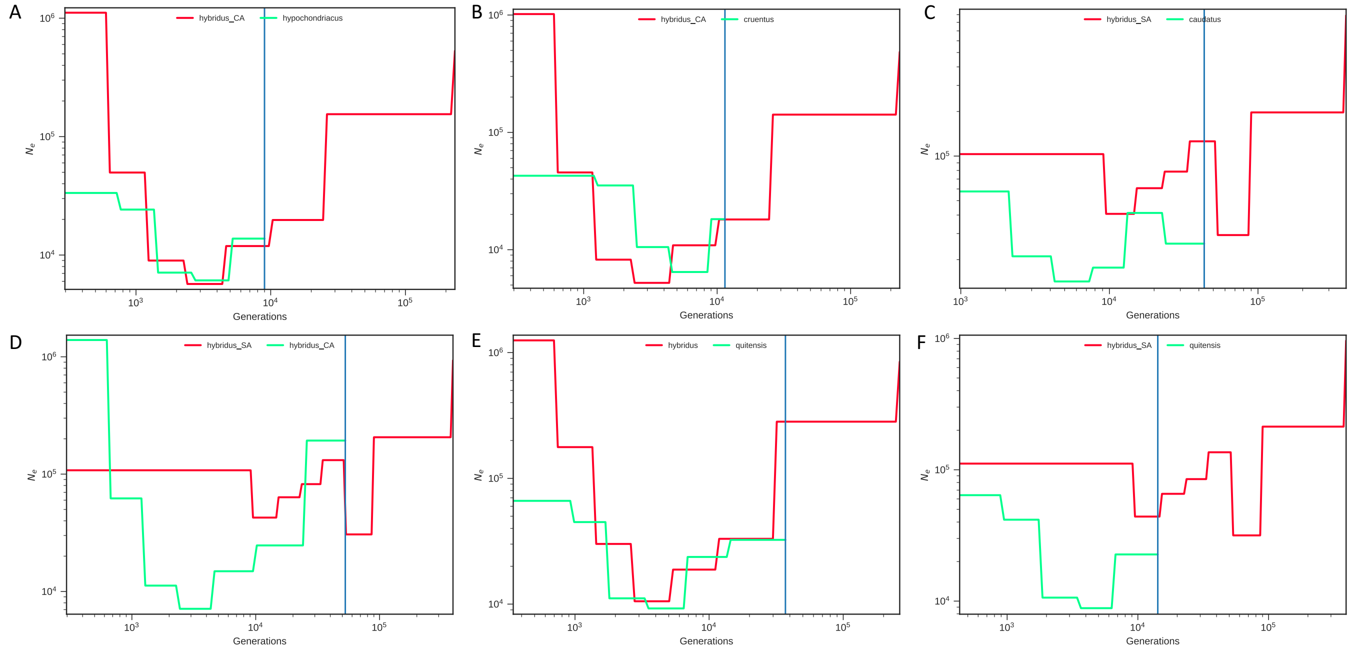


**Figure S19** Split times of crops species from sepatate hybridus (SA= South America; CA = Central America) populations.

### Supplementary Tables

**Table S1** Accessions sequenced in this study and seed color denominated from the same seed bag as sequenced individuals. Accessions denoted as ′not clear′ had seeds of both seed colors in the seed lot distributed by the USDA germplasm collection and were removed from the GWAS analysis

| No | Accession number | Species | Origcty | Seed color |
|---|---|---|---|---|
| 1 | PI 490491 | caudatus | ARG | dark |
| 2 | PI 511712 | caudatus | ECU | dark |
| 3 | PI 608019 | caudatus | ECU | dark |
| 4 | PI 649228 | caudatus | PER | dark |
| 5 | PI 490511 | caudatus | PER | dark |
| 6 | PI 490518 | caudatus | PER | dark |
| 7 | PI 481949 | caudatus | PER | dark |
| 8 | PI 511686 | caudatus | PER | dark |
| 9 | PI 649217 | caudatus | PER | not clear |
| 10 | PI 511690 | caudatus | PER | not clear |
| 11 | PI 511680 | caudatus | ARG | white |
| 12 | PI 511679 | caudatus | ARG | white |
| 13 | PI 490459 | caudatus | BOL | white |
| 14 | PI 511681 | caudatus | BOL | white |
| 15 | PI 490604 | caudatus | BOL | white |
| 16 | PI 642741 | caudatus | BOL | white |
| 17 | AMA 107 | caudatus | CHN | white |
| 18 | PI 490609 | caudatus | ECU | white |
| 19 | PI 649230 | caudatus | PER | white |
| 20 | PI 511704 | caudatus | PER | white |
| 21 | PI 649227 | caudatus | PER | white |
| 22 | AMA 125 | caudatus | PER | white |
| 23 | PI 481957 | caudatus | PER | white |
| 24 | PI 511706 | caudatus | PER | white |
| 25 | PI 490561 | caudatus | PER | white |
| 26 | PI 511687 | caudatus | PER | white |
| 27 | PI 490612 | caudatus | PER | white |
| 28 | PI 490431 | caudatus | PER | white |
| 29 | PI 511696 | caudatus | PER | white |
| 30 | PI 481965 | caudatus | PER | white |
| 31 | Ames 5302 | caudatus | PER | white |
| 32 | PI 481960 | caudatus | PER | white |
| 33 | PI 511717 | cruentus | GTM | dark |
| 34 | PI 667160 | cruentus | GTM | dark |
| 35 | PI 658728 | cruentus | MEX | dark |
| 36 | PI 511714 | cruentus | PER | dark |
| 37 | PI 511713 | cruentus | PER | dark |
| 38 | AMA 155 | cruentus | USA | dark |
| 39 | PI 667165 | cruentus | BRA | not clear |
| 40 | PI 643049 | cruentus | MEX | not clear |
| 41 | Baernkrafft | cruentus | DEU | white |
| 42 | PI 433228 | cruentus | GTM | white |
| 43 | PI 451826 | cruentus | GTM | white |
| 44 | PI 643039 | cruentus | MEX | white |
| 45 | PI 643058 | cruentus | MEX | white |
| 46 | PI 649514 | cruentus | MEX | white |
| 47 | PI 606798 | cruentus | MEX | white |
| 48 | PI 576482 | cruentus | MEX | white |
| 49 | PI 576481 | cruentus | MEX | white |
| 50 | PI 649609 | cruentus | MEX | white |
| 51 | PI 511723 | cruentus | MEX | white |
| 52 | PI 511876 | cruentus | MEX | white |
| 53 | Ames 5552 | cruentus | MEX | white |
| 54 | PI 643042 | cruentus | MEX | white |
| 55 | PI 649524 | cruentus | MEX | white |
| 56 | PI 643037 | cruentus | MEX | white |
| 57 | PI 649509 | cruentus | MEX | white |
| 58 | PI 604571 | hybr. | MEX | dark |
| 59 | PI 636180 | hybridus | COL | dark |
| 60 | PI 490684 | hybridus | ECU | dark |
| 61 | PI 511754 | hybridus | ECU | dark |
| 62 | PI 490679 | hybridus | ECU | dark |
| 63 | PI 490689 | hybridus | ECU | dark |
| 64 | PI 490670 | hybridus | ECU | dark |
| 65 | PI 490664 | hybridus | ECU | dark |
| 66 | PI 490739 | hybridus | ECU | dark |
| 67 | PI 667156 | hybridus | ECU | dark |
| 68 | PI 490731 | hybridus | ECU | dark |
| 69 | PI 667158 | hybridus | GTM | dark |
| 70 | PI 604582 | hybridus | MEX | dark |
| 71 | PI 604568 | hybridus | MEX | dark |
| 72 | PI 604574 | hybridus | MEX | dark |
| 73 | PI 511724 | hybridus | MEX | dark |
| 74 | Ames 5232 | hybridus | PER | dark |
| 75 | PI 490740 | hybridus | PER | dark |
| 76 | PI 490489 | hybridus | PER | dark |
| 77 | Ames 5335 | hybridus | BOL | not clear |
| 78 | PI 652432 | hypochondriacus | BRA | dark |
| 79 | PI 604581 | hypochondriacus | MEX | dark |
| 80 | PI 649602 | hypochondriacus | MEX | dark |
| 81 | PI 604587 | hypochondriacus | MEX | dark |
| 82 | Ames 2085 | hypochondriacus | MEX | not clear |
| 83 | PI 649607 | hypochondriacus | MEX | white |
| 84 | PI 649587 | hypochondriacus | MEX | white |
| 85 | PI 649537 | hypochondriacus | MEX | white |
| 86 | PI 643036 | hypochondriacus | MEX | white |
| 87 | PI 649575 | hypochondriacus | MEX | white |
| 88 | Ames 5457 | hypochondriacus | MEX | white |
| 89 | PI 633589 | hypochondriacus | MEX | white |
| 90 | PI 649565 | hypochondriacus | MEX | white |
| 91 | PI 604559 | hypochondriacus | MEX | white |
| 92 | PI 643070 | hypochondriacus | MEX | white |
| 93 | PI 643041 | hypochondriacus | MEX | white |
| 94 | PI 604595 | hypochondriacus | MEX | white |
| 95 | PI 649529 | hypochondriacus | MEX | white |
| 96 | Ames 2215 | hypochondriacus | MEX | white |
| 97 | PI 511731 | hypochondriacus | MEX | white |
| 98 | PI 643067 | hypochondriacus | MEX | white |
| 99 | PI 649559 | hypochondriacus | MEX | white |
| 100 | PI 649595 | hypochondriacus | MEX | white |
| 101 | PI 649623 | hypochondriacus | MEX | white |
| 102 | AMA 89 | powellii | RWA | dark |
| 103 | Ames 21666 | quitensis | ARG | dark |
| 104 | Ames 5334 | quitensis | ARG | dark |
| 105 | PI 511736 | quitensis | BOL | dark |
| 106 | PI 652426 | quitensis | BRA | dark |
| 107 | PI 652428 | quitensis | BRA | dark |
| 108 | PI 652422 | quitensis | BRA | dark |
| 109 | PI 511747 | quitensis | ECU | dark |
| 110 | PI 511749 | quitensis | ECU | dark |
| 111 | PI 511737 | quitensis | ECU | dark |
| 112 | PI 490705 | quitensis | ECU | dark |
| 113 | PI 511745 | quitensis | ECU | dark |
| 114 | PI 490720 | quitensis | ECU | dark |
| 115 | PI 511741 | quitensis | ECU | dark |
| 116 | Ames 5247 | quitensis | ECU | dark |
| 117 | PI 490673 | quitensis | ECU | dark |
| 118 | PI 649246 | quitensis | PER | dark |
| 119 | PI 511751 | quitensis | PER | dark |
| 120 | Ames 5342 | quitensis | PER | dark |
| 121 | PI 490466 | quitensis | PER | dark |

**Table S2** Proportion of mapped reads by species.

| Species | mapped reads (%) | standard deviation |
|---|---|---|
| caudatus | 94.81594 | 8.720722 |
| cruentus | 91.98360 | 16.291917 |
| hypochondriacus | 94.35625 | 14.640438 |
| hybridus | 92.06421 | 16.640896 |
| quitensis | 96.56000 | 1.438093 |
| hybr. | 96.98000 | NA |
| powellii | 96.70000 | NA |

**Table S3** Candidate genes for seed color in grain amaranth

| Chromosome | Start | End | Name | Strand | Description |
|---|---|---|---|---|---|
| 3 | 5881516 | 5896384 | AH005257 | + | Similar to CCT1: T-complex protein 1 subunit alpha (Arabidopsis thaliana) |
| 3 | 5897584 | 5905365 | AH005258 | + | Similar to MLO13: MLO-like protein 13 (Arabidopsis thaliana) |
| 3 | 5906387 | 5930826 | AH005259 | - | Similar to NUP160: Nuclear pore complex protein NUP160 (Arabidopsis thaliana) |
| 3 | 5933604 | 5946653 | AH005260 | - | Similar to CMO: Choline monooxygenase, chloroplastic (Amaranthus tricolor) |
| 3 | 5959962 | 5966304 | AH005261 | - | Similar to Ank3: Ankyrin-3 (Mus musculus) |
| 9 | 14260388 | 14267067 | AH014529 | - | Similar to NYC1: Probable chlorophyll(ide) b reductase NYC1, chloroplastic (Oryza sativa subsp. japonica) |
| 9 | 14276572 | 14283316 | AH014530 | + | Protein of unknown function |
| 9 | 14291900 | 14298118 | AH014531 | - | Protein of unknown function |
| 9 | 14298147 | 14298790 | AH014532 | - | Protein of unknown function |
| 9 | 14298908 | 14299399 | AH014533 | - | Protein of unknown function |
| 9 | 14304669 | 14312794 | AH014534 | + | Similar to CSLG2: Cellulose synthase-like protein G2 (Arabidopsis thaliana) |
| 9 | 14333735 | 14334892 | AH014535 | + | Similar to rmnd5a: Protein RMD5 homolog A (Xenopus laevis) |
| 9 | 14345025 | 14345715 | AH014536 | + | Similar to OFP8: Transcription repressor OFP8 (Arabidopsis thaliana) |
| 9 | 14353195 | 14353923 | AH014537 | + | Protein of unknown function |
| 9 | 14355443 | 14356829 | AH014538 | - | Similar to TRX2: Thioredoxin H2 (Arabidopsis thaliana) |
| 9 | 14366001 | 14367256 | AH014539 | - | Similar to TRX2: Thioredoxin H2 (Arabidopsis thaliana) |
| 9 | 14372147 | 14375412 | AH014540 | - | Similar to HSFC1: Heat stress transcription factor C-1 (Arabidopsis thaliana) |
| 9 | 14378777 | 14389557 | AH014541 | - | Similar to KIN13B: Kinesin-like protein KIN-13B (Arabidopsis thaliana) |
| 9 | 14396256 | 14396438 | AH014542 | + | Protein of unknown function |
| 9 | 14404615 | 14409777 | AH014543 | - | Similar to Os01g0583100: Probable protein phosphatase 2C 6 (Oryza sativa subsp. japonica) |
| 9 | 14412914 | 14418029 | AH014544 | - | Similar to CFIS1: Pre-mRNA cleavage factor Im 25 kDa subunit 1 (Arabidopsis thaliana) |
| 9 | 14420597 | 14423542 | AH014545 | - | Similar to At4g06634: Uncharacterized zinc finger protein At4g06634 (Arabidopsis thaliana) |
| 9 | 14443999 | 14450734 | AH014546 | + | Protein of unknown function |
| 9 | 14467556 | 14471512 | AH014547 | + | Protein of unknown function |
| 9 | 14472750 | 14477103 | AH014548 | - | Similar to UBA1C: UBP1-associated proteins 1C (Arabidopsis thaliana) |
| 9 | 14480031 | 14487685 | AH014549 | + | Protein of unknown function |
| 9 | 14503360 | 14505015 | AH014550 | + | Protein of unknown function |
| 9 | 14513815 | 14518528 | AH014551 | + | Protein of unknown function |
| 9 | 14525887 | 14543181 | AH014552 | + | Similar to XI-K: Myosin-17 (Arabidopsis thaliana) |
| 9 | 14545885 | 14551450 | AH014553 | - | Similar to SPBC776.05: Uncharacterized membrane protein C776.05 (Schizosaccharomyces pombe (strain 972 / ATCC 24843)) |
| 9 | 14552847 | 14558739 | AH014554 | - | Similar to AL5: PHD finger protein ALFIN-LIKE 5 (Arabidopsis thaliana) |
| 9 | 14580201 | 14584739 | AH014555 | + | Protein of unknown function |
| 9 | 14587900 | 14593548 | AH014556 | + | Similar to NFYA6: Nuclear transcription factor Y subunit A-6 (Arabidopsis thaliana) |
| 9 | 14598265 | 14609302 | AH014557 | + | Similar to BRXL4: Protein Brevis radix-like 4 (Arabidopsis thaliana) |
| 9 | 14610048 | 14614953 | AH014558 | + | Similar to NLP3: Omega-amidase, chloroplastic (Arabidopsis thaliana) |
| 9 | 14622402 | 14624168 | AH014559 | + | Similar to At1g54200: Protein BIG GRAIN 1-like B (Arabidopsis thaliana) |
| 9 | 14641501 | 14645682 | AH014560 | + | Similar to VPS2.1: Vacuolar protein sorting-associated protein 2 homolog 1 (Arabidopsis thaliana) |
| 9 | 14650590 | 14651111 | AH014561 | - | Similar to ATL53: Putative RING-H2 finger protein ATL53 (Arabidopsis thaliana) |
| 9 | 14657567 | 14664508 | AH014562 | + | Similar to MSL10: Mechanosensitive ion channel protein 10 (Arabidopsis thaliana) |
| 9 | 14665596 | 14668727 | AH014563 | - | Similar to RPA1A: Replication protein A 70 kDa DNA-binding subunit A (Arabidopsis thaliana) |
| 9 | 14677913 | 14679912 | AH014564 | - | Similar to EFL4: Protein ELF4-LIKE 4 (Arabidopsis thaliana) |
| 9 | 14700127 | 14700888 | AH014565 | + | Similar to UP3: Stress-response A/B barrel domain-containing protein UP3 (Arabidopsis thaliana) |
| 9 | 14705869 | 14708414 | AH014566 | + | Similar to C1: Anthocyanin regulatory C1 protein (Zea mays) |
| 9 | 14713047 | 14716244 | AH014567 | - | Similar to RPS3A: 40S ribosomal protein S3-1 (Arabidopsis thaliana) |
| 9 | 14722269 | 14730995 | AH014568 | + | Protein of unknown function |
| 9 | 14734261 | 14737531 | AH014569 | - | Similar to GRF5: Growth-regulating factor 5 (Arabidopsis thaliana) |
| 9 | 14759660 | 14776842 | AH014570 | - | Protein of unknown function |
| 9 | 14780031 | 14782777 | AH014571 | + | Similar to OPR3: 12-oxophytodienoate reductase 3 (Solanum lycopersicum) |
| 9 | 14783927 | 14788106 | AH014572 | - | Similar to REM16: B3 domain-containing protein REM16 (Arabidopsis thaliana) |
| 9 | 14792503 | 14796787 | AH014573 | - | Similar to REM10: B3 domain-containing protein REM10 (Arabidopsis thaliana) |
| 9 | 14805984 | 14809221 | AH014574 | + | Similar to OPR3: 12-oxophytodienoate reductase 3 (Solanum lycopersicum) |
| 9 | 14814160 | 14814522 | AH014575 | - | Similar to NFYB6: Nuclear transcription factor Y subunit B-6 (Arabidopsis thaliana) |
| 9 | 14830784 | 14835140 | AH014576 | - | Similar to GALS1: Galactan beta-1,4-galactosyltransferase GALS1 (Arabidopsis thaliana) |
| 9 | 14842490 | 14846361 | AH014577 | - | Similar to AMC5: Metacaspase-5 (Arabidopsis thaliana) |
| 9 | 14855046 | 14855666 | AH014578 | - | Similar to ABP19A: Auxin-binding protein ABP19a (Prunus persica) |

### Supplementary Notes

#### *Species delimitation*

For six accessions, the species assignment in the genebank passport data did not match the results of the genetic analysis (Table S4) and may either represent misclassified accessions or mixup of seeds during seed handling and regeneration. These should be reconsidered for morphological classification. A possible misclassification is of particular interest interesting for the only amaranth variety ever registered in Germany, which considered to be *A. cruentus*, but according to the genetic data appears to be *A. hypochondriacus*.

**Table S4** Accessions in which the species assignment in the genebank passport data is not supported by the genetic analysis.

| Individual | Genebank annotation | Genomic group |
|---|---|---|
| PI604587 | *A. hypochondriacus* | *A. cruentus* |
| Ames2215 | *A. hypochondriacus* | *A. cruentus* |
| PI604595 | *A. hypochondriacus* | *A. cruentus* |
| Baernkrafft (German variety) | *A. cruentus* | *A. hypochondriacus* |
| PI511723 | *A. cruentus* | *A. hypochondriacus* |
| PI511876 | *A. cruentus* | *A. hypochondriacus* |
